# MAGELLAN: Automated Generation of Interpretable Computational Models for Biological Reasoning

**DOI:** 10.1101/2025.05.16.653408

**Authors:** Matthew A. Clarke, Charlie George Barker, Yuxin Sun, Theodoros I. Roumeliotis, Jyoti S. Choudhary, Jasmin Fisher

**Affiliations:** UCL Cancer Institute, University College London, 72 Huntley Street, London, WC1E 6DD, UK; The Institute of Cancer Research, 123 Old Brompton Road, London, SW7 3RP, UK

## Abstract

Computational models have become essential tools for understanding signalling networks and their non-linear dynamics. However, these models are typically constructed manually using prior knowledge and can be over-reliant on study bias. These limitations hinder their ability to make accurate predictions and incorporate new evidence. Scaling up the construction of models to take advantage of increasingly abundant ‘omics data can bridge these gaps by providing a comprehensive view of signalling events and how they influence cellular phenotypes. In this study, we present MAGELLAN, a method leveraging message passing graph neural networks to build computational models directly from pathway data and discrete rules representing experimental results. We used this to construct a computational model of breast cancer signalling and re-parameterize a previously published non-small cell lung cancer (NSCLC) model, showing that MAGELLAN can predict genetic dependencies and achieve comparable model quality to expert-curated and manually trained models. Our approach enables the integration of prior knowledge networks and experimental data to build predictive models that are mechanistically interpretable. This approach simplifies model creation, making it more accessible and practical for experimentalists, and supports broader applications in drug discovery and biological research.

## Introduction

Signalling proteins play important roles in the development of cancer, frequently harbouring mutations that drive oncogenesis and therapeutic resistance^1^. Computational modelling offers a powerful tool to predict molecular signalling processes and the resultant biological behaviours (such as cell growth and survival) by efficiently simulating networks of interacting biological species and proteins. These proteins often exhibit a sigmoidal activation profile and can therefore be abstracted into discrete entities^2,3^, which can be used to efficiently represent their activity and simulate their dynamics. To be useful for drug design and advancing our understanding of fundamental biological mechanisms, these simulations should be mechanistically interpretable. However, the manual construction of mechanistically interpretable models of biological processes and the validation of these complex models is laborious and time-consuming, requiring extensive biological expertise and iterative refinement. Furthermore, the nature of this biological expertise tends to focus disproportionately on highly cited proteins, driven by conservative study biases^4,5^.

Large scale ‘omics technologies have been used to explore new dimensions of signalling with notable success^6–11^, and the rate at which new data is generated is increasing. These data are suited to analysis with machine learning, which can handle large feature sets effectively^8,12,13^. However, this approach comes with a caveat: the sacrifice of interpretability in favour of predictive accuracy. Mechanistic insight and accessibility are essential concepts for translatable research, as they offer transparency to clinicians and support hypothesis-driven experimentation.

Executable models are a well-developed paradigm of computational models whose utility has been consistently proven^14–20^. They treat biological systems as networks whose interactions can be thought of as biological ‘machines’ executing a program encoded in the structure of the network. Qualitative networks^21^ are an extension of Boolean networks^22^, which can be ‘executed’ by allowing nodes to interact dynamically with one another until a stable state of the network is reached, which can be interpreted as an overall cellular phenotype^21^. This enables the creation of mechanistic models of complex biological systems that can be used to understand emergent properties derived from prior knowledge^23^. Although these models are computationally efficient, constructing these models is manual, making them laborious to do at scale. We posit that combining the mechanistic interpretability of executable models^15^ with the data-driven power of machine learning will enable a deeper understanding of signalling pathways and allow the development of more novel and effective therapeutic strategies in an accessible and transparent manner.

To address this, we present the optimisation tool, MAGELLAN (**M**essage p**A**ssing **G**raph n**E**uraL biomode**LA**nalyzer **N**etwork), which uses a machine-learning-based parameterization approach for semi-automatically building and testing models which can then be further explored and built upon using the open-source BioModelAnalyzer (BMA) tool (https://biomodelanalyzer.org/). We benchmark this algorithm on simulated contexts and a previously published computational model of non-small cell lung cancer (NSCLC)^24^. We then applied MAGELLAN to construct a computational model of breast cancer tumour intrinsic signalling based on a literature curated specification. These qualitative models are capable of replicating key behaviours observed in the training dataset and generating novel predictions of essentiality data. Our method was developed with a focus on interpretability and accessibility, aiming to serve as a ‘co-pilot’ for experimentalists by suggesting targeted experiments to fill gaps in the model.

This approach helps researchers identify the most prudent next experiments in their investigations based on previously supplied information. It then iteratively builds models that illuminate the mechanisms underlying experimental findings in a semi-automated fashion. MAGELLAN is compatible with the existing graphical user interface (GUI) of the BMA tool, allowing models to be loaded, simulated, and edited through ‘point-and-click’ functionality within a web browser. This makes the tool accessible to researchers who may not have experience with coding and command-line tools.

In this study, we show how MAGELLAN can be used to explore signalling pathways necessary for cell growth in various contexts. By integrating sparse, qualitative behavioural observations, taken from the literature, with prior-knowledge networks, we uncover drivers of oncogenesis that may be targeted for therapeutic intervention. MAGELLAN bridges the data-driven nature of machine learning approaches with the interpretive power of computational modelling. This enables researchers to predict biological processes and place their findings within a broader mechanistic framework, ultimately facilitating the discovery of therapeutic targets and guiding precision medicine efforts.

## Results

### MAGELLAN: a tool for automated generation of executable biological models

MAGELLAN is an optimisation algorithm for qualitative networks using interpretable message-passing graph neural networks (GNNs). It uses user-specified prior knowledge graphs to construct executable biological models given a set of known experimental behaviours. These models are computer programs whose execution simulates a cell’s behaviour^25^, allowing for mechanistically interpretable predictions of biological processes^21,23^. We use the GNN to fit the edge weights of the model such that it can recapitulate a set of discrete behaviours taken from varied experimental or literature-based provenance. These specifications define qualitative relationships between initial conditions, such as genetics or environmental perturbations, and resultant biological outcomes, encompassing both high-level phenotypic traits (e.g., apoptosis, proliferation) and low-level molecular activities (e.g., kinase activities). Here, we use MAGELLAN to fit such qualitative specifications, benchmarking on various contexts (Figure 1A).

**Figure 1.**
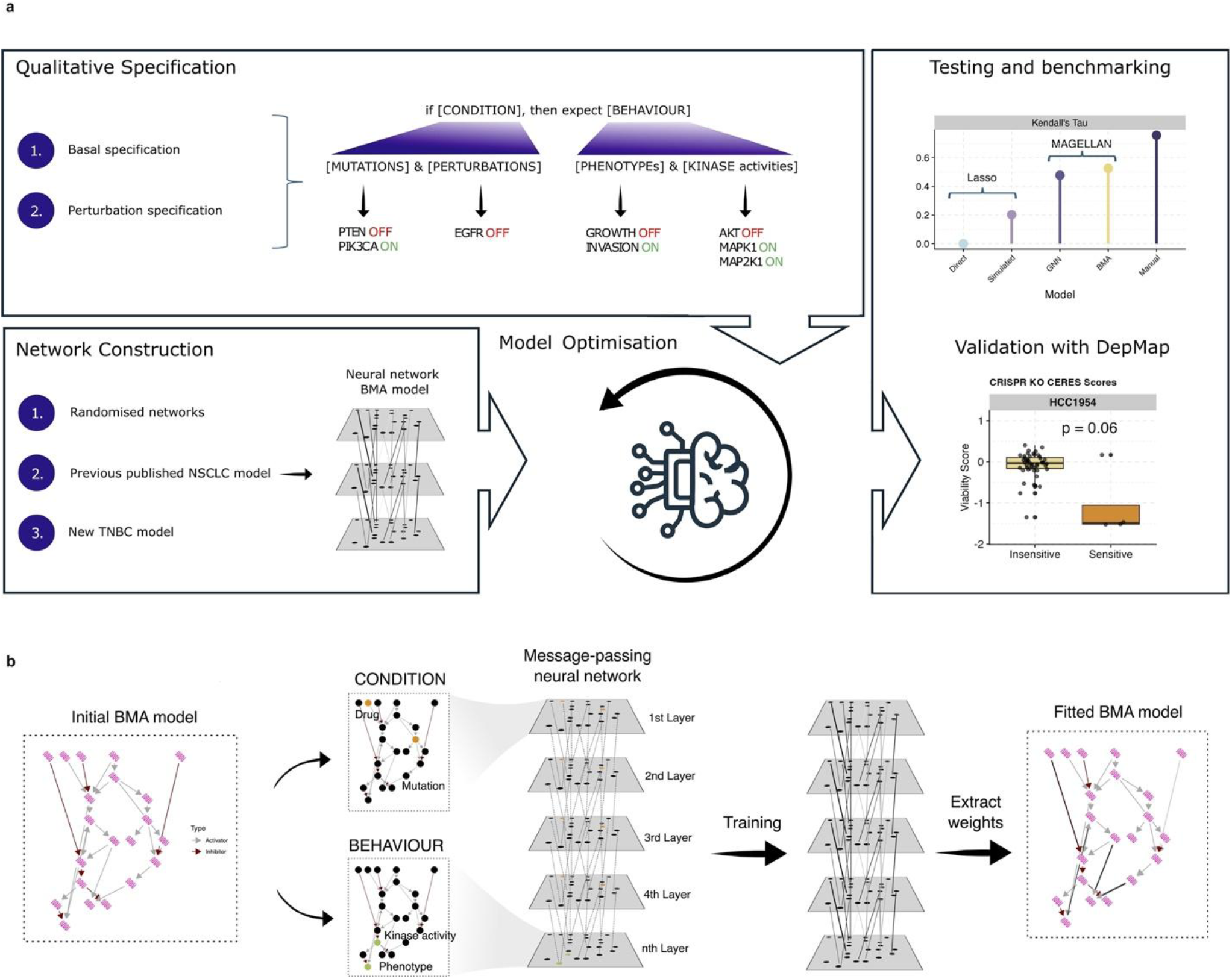
Outline of the workflow and basic fitting of a computational model to the literature derived specification using MAGELLAN. **A)** Schematic illustrating the workflow of the paper, as well as the composition and structure of the specification used to train the models. **B)** Schematic showing how MAGELLAN translates biological interaction networks (left) into a neural network to fit weights to behaviours (right), given sets of conditions. For more information see Methods.

The methodology of MAGELLAN involves the transformation of a prior knowledge network into a GNN architecture. Nodes within this network represent biological species with discrete biological activities, while directed edges signify regulatory interactions between them. The GNN’s layered structure mirrors the temporal propagation of activities within the biological system, with each consecutive layer corresponding to a timepoint in the simulation (see Methods). Initial conditions defined by the specification are encoded as input states, and the GNN is trained to predict corresponding output behaviours (see Methods). Iterative optimization of edge weights refines the model, emphasizing interactions critical for translating input conditions into observed phenotypes, while minimizing the influence of irrelevant or noisy edges. The resulting weighted network constitutes an executable model, which can be efficiently deployed for simulation and analysis using the BioModelAnalyzer (BMA) webtool. This approach enables the derivation of mechanistically interpretable insights into biological network dynamics, bridging prior knowledge with empirical observations (Figure 1B).

### MAGELLAN is scalable and robust when tested on synthetic benchmarks

We evaluated the performance of MAGELLAN by comparing the behaviour of a reconstructed model, obtained through the optimisation process, with that of the original breast cancer model for synthetic specifications. This comparison was based on a subset of experiments from the specification that were set aside during training (see Methods). This comparison allowed us to quantify MAGELLAN’s ability to accurately infer model parameters from simulated biological data, thereby demonstrating its potential for application to complex, real-world biological systems. We tested the model for different sizes of network (Figure 2A) and found that it scales well to larger sizes (1000s of edges). We further tested the ability to correctly recover behaviour that matches the specification and found that this is robust for both small specifications compared to the number of nodes in the network (Supplementary Figure 1A).

**Figure 2.**
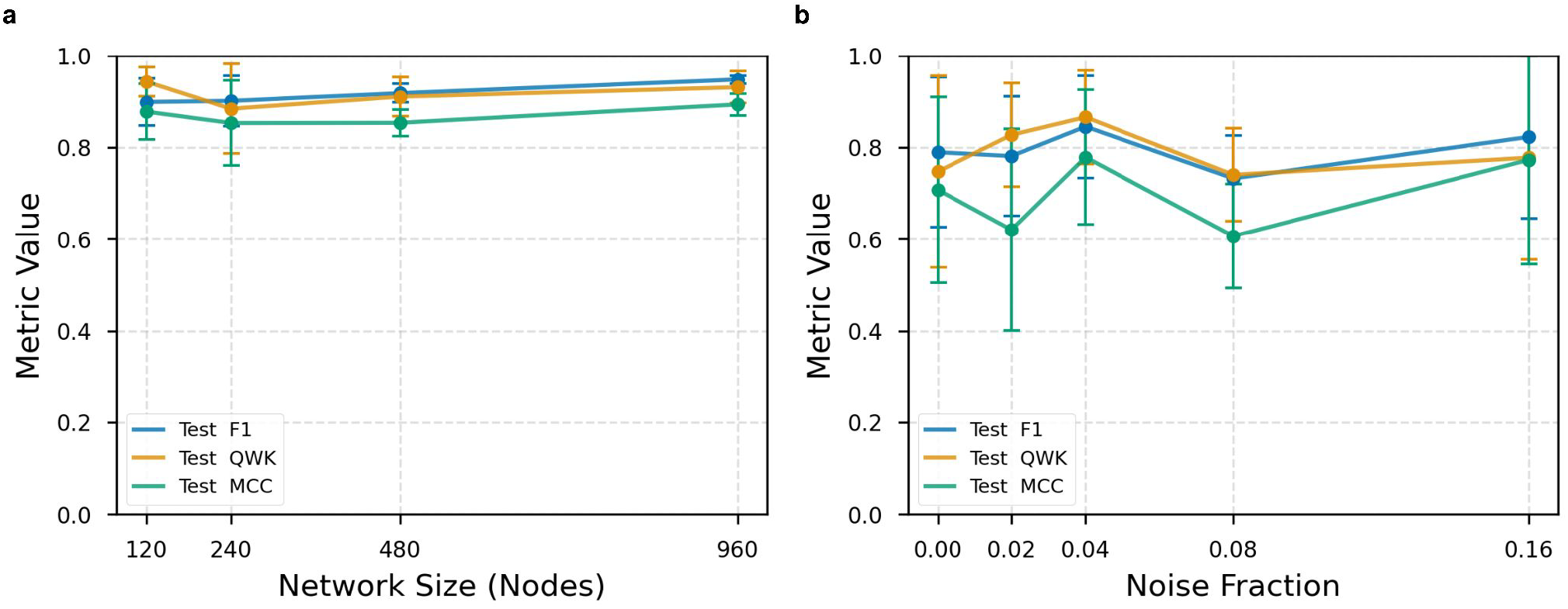
Using synthetic benchmarks to test MAGELLAN’s ability to recover network dynamics. **A)** Effect of network size (x axis) on the quality of fitting to a test set (y axis). **B)** Effect of including noisy or incorrect edges (noise fraction, x axis) on the quality of fitting to the testing set (y axis). Model fitting is evaluated by various metrics (F1, MCC, QWK) for a 10% test set of the specification (see Methods).

Real biological data contain noise and errors, so we assessed MAGELLANs ability to handle inaccuracies in the specification used for fitting. We found that with 10% of the measurements per experiment in the specification being incorrect, the quality of the fitting remains largely unaffected (Figure 2B). Finally, we assessed MAGELLAN’s ability to eliminate spurious edges while retaining true ones and found that MAGELLAN exhibited a low false-positive rate but was conservative in its pruning. This is due to the need for a large coverage of low-level measured nodes to be able to distinguish between different redundant pathways through the network (Supplementary Figure 1B, C). However, as a specification incorporates experiments measuring more nodes in the network, the number of false edges removed increases while the rate of true edges removed remains low (Supplementary Figure 1B). Satisfied that MAGELLAN can accurately recover ground truth synthetic model behaviours, we moved to more complex, real-world biological contexts.

### Automated parameterization with MAGELLAN replicates manual model tuning in minutes

We applied MAGELLAN to a previously published 160-node, 360-edge NSCLC computational model^24^. This model, encompassing key oncogenic pathways, was manually fitted to a specification derived from 26 publications over 12 months and validated by further experiments. The specification detailed rates of growth, apoptosis, and protein activation from varied experimental conditions. This extensive manual effort provided a rigorous benchmark for MAGELLAN’s automated fitting capabilities against real reasoning from experts in the field. Cell lines were modelled by setting nodes corresponding to genes with known driver mutations to fixed values (0 for loss-of-function mutations and 3 for gain-of-function mutations). This was then combined with initialisations representing the conditions in the experiments tested. Mutation data for these cell lines were obtained from Cell Model Passports^26^ (see Methods).

We further benchmarked MAGELLAN and manual parameterization against a naive approach using iterative Lasso regression, which employs linear modelling with L1 regularization to fit network weights (see Methods). The results of the fitting can be visualised and simulated using the web-based BMA tool (https://biomodelanalyzer.org/). We found that MAGELLAN is relatively strict at removing edges not necessary for model functioning, removing 12 edges preserved in manual training (Supplementary Table S1). This MAGELLAN-optimised network (Figure 3A, Supplementary Figure 2) outperforms lasso regression-based techniques (Quadratic Weighted Kappa, Figure 3B) at fitting the specification (Supplementary Table S2). Notably, MAGELLAN achieved QWK scores comparable to those obtained through manual parameter optimization. The manual training represents an extensive effort involving domain experts refining model parameters over several months. In contrast, MAGELLAN achieved comparable performance within two minutes of computational processing.

**Figure 3.**
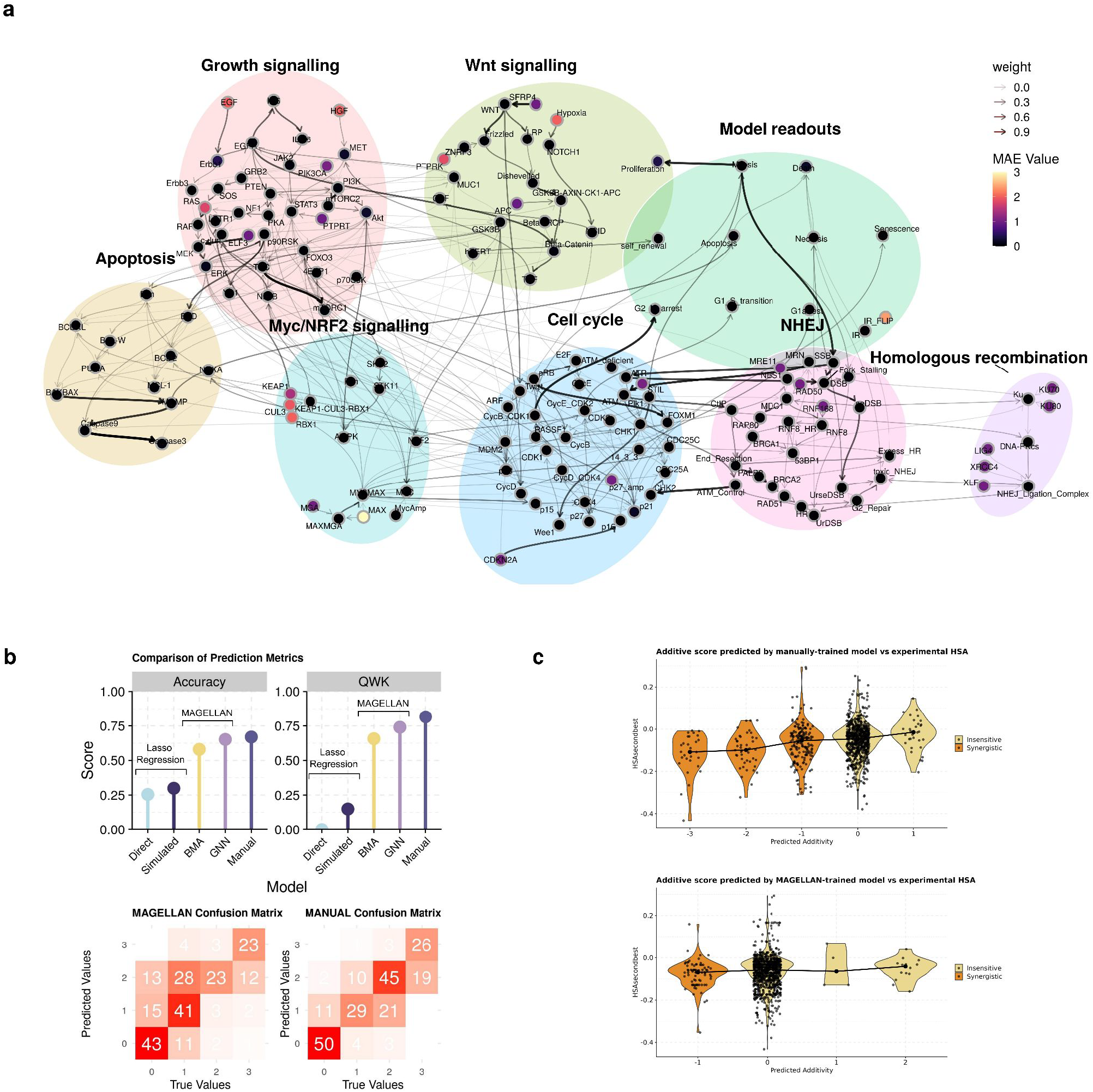
Benchmarking of MAGELLAN using a NSCLC computational model. **A)** Network visualization depicting the results of model fitting. Nodes are coloured according to their Mean Absolute Error (MAE). Edge colour and intensity represent the fitted edge weights. Edges represent regulatory interactions between proteins and are pointed for activating edges (→) and blunt for inhibiting (⊣). **B)** Assessment of model fit against manually parameterised model from Clarke *et al*. (2025)^24^, showing accuracy, Kendall’s Tau and Quadratic Weighted Kappa for different methods of fitting the specification to the network. **C)** Violin plots showing the additive score (x axis, see Methods) as predicted by the manually trained (top) and MAGELLAN-trained (bottom) model versus the HSA measured experimentally by Nair et al. (2022) (y axis). Predicted additivity below 0 (representing synergistic combinations) are coloured orange and additive scores 0 and above (representing insensitive combinations) are coloured pale yellow.

To assess the utility of the NSCLC model, we compared the predictions of experimentally tested combination therapy screens to the predictions of manually and MAGELLAN-trained NSCLC models. This assessed the additive effect of combining various drug treatments in six NSCLC cell models, SW1573, NCIH1792, HCC827, PC9, A549 and NCIH1993. Additive score is calculated for each drug combination and measures the overall survival (Proliferation – Death) of drug treatments when compared to the most effective single treatment (see Methods). We compared this to the Highest Single Agent effect (HSA) as measured by Nair *et al*. (2022)^27^ for the cell lines tested (Figure 3C). The manually trained model is more granular in its predictions of growth, with more combinations predicted to have an effect on growth and with a greater number of increments. The MAGELLAN trained model is more conservative with its predictions, reflecting our observations from the simulated data. Nonetheless, when comparing the HSA additivity score between additive and insensitive predictions from both models we see significant changes. For the manually trained NSCLC model, we observe that predicted synergistic combinations have significantly lower HSA values than those predicted to be insensitive (Welch’s two-sample t-tests, p = 1.788 × 10^−5^). For the MAGELLAN-predicted drug combinations, we observe a marginally significant trend where predicted synergistic combinations have lower HSA values than those predicted to be insensitive (Welch’s two-sample t-tests, p = 0.09). These results indicate that both MAGELLAN and the manually trained model are able to differentiate between synergistic and insensitive drug combinations. However, the manually-trained model shows a greater degree of statistical significance and both MAGELLAN and the manually-trained model are conservative in their prediction of therapies that interact in an additive fashion.

### Using MAGELLAN to train an explainable model of breast cancer signalling with prior-knowledge data

Beyond benchmarking MAGELLAN on pre-optimized networks, we aimed to examine its utility in automatically generated networks from prior-knowledge databases. This context makes MAGELLAN more accessible to users without *a priori* experience in constructing models. To illustrate the utility of MAGELLAN fitting a prior-knowledge based network, we used it to train a basic computational model of breast cancer growth signalling. We automatically constructed a breast cancer signalling network using the pathway-database KEGG^28^, combining the ‘Breast Cancer Pathway’ (HSA05224) and the ‘Estrogen signalling pathway’ (HSA04915) and translating this network into a BMA executable model (see Methods). These pathways were selected because they contain the main regulatory processes associated with breast cancer oncogenesis.

The resultant network contains nodes to represent each protein, with a corresponding target function defining its activity. This activity is described by a discrete value between 3 states. A state of ‘0’ indicates reduced activity, ‘1’ basal activity and ‘2’ elevated activities in a given context. Note, that the number of states in the qualitative network model is a user-defined hyperparameter and can be adjusted to suit the granularity of the training specification. This produces a model of 60 nodes, representing proteins and biological entities and 93 regulatory edges between them (Supplementary Table S3).

We first compiled a set of behaviours without perturbation in 7 breast cell lines, MCF10A, MCF7, MDA-MB-231, T47D, BT474, MDA-MB-468 and HCC1954 (Supplementary Table S4). This specification includes simple behaviours, such as ‘KRAS mutation leads to growth,’ ‘TNFα stimulation induces cell death,’ or ‘EGFR stimulation activates MAPK.’ Next, we compiled a specification dataset describing experiments derived from a curated list of 10 publications. These cover stimulation experiments with specific growth factors^29–32^, perturbation experiments^33–36^ with common breast cancer therapies and overexpression to transform otherwise non-tumorigenic cell lines into tumorigenic breast cancer^37,38^. These rules constrain the model’s behaviour and predictions to ground truths taken from the literature. We combined these conditions with the genetics of the cell lines from Cell Model Passports^26^ (Supplementary Table S5) to initialise the model for different experiments.

Using MAGELLAN, we automatically fit the weights of the model (Figure 4A), showing that it reproduces the main features of the specification set (Figure 4B-C, Supplementary Table S4). Through the progress of this training, initial fits would remove the MAPK pathway upstream of KRAS. Accordingly, we searched for experiments in the literature for perturbations to MAPK to add to the literature-derived specification^39^. In this way, we illustrate the semi-automated nature of MAGELLAN and how it can be used to guide experimental biologists to the optimal experiments required to bridge knowledge-gaps represented in the model and from current training data.

**Figure 4.**
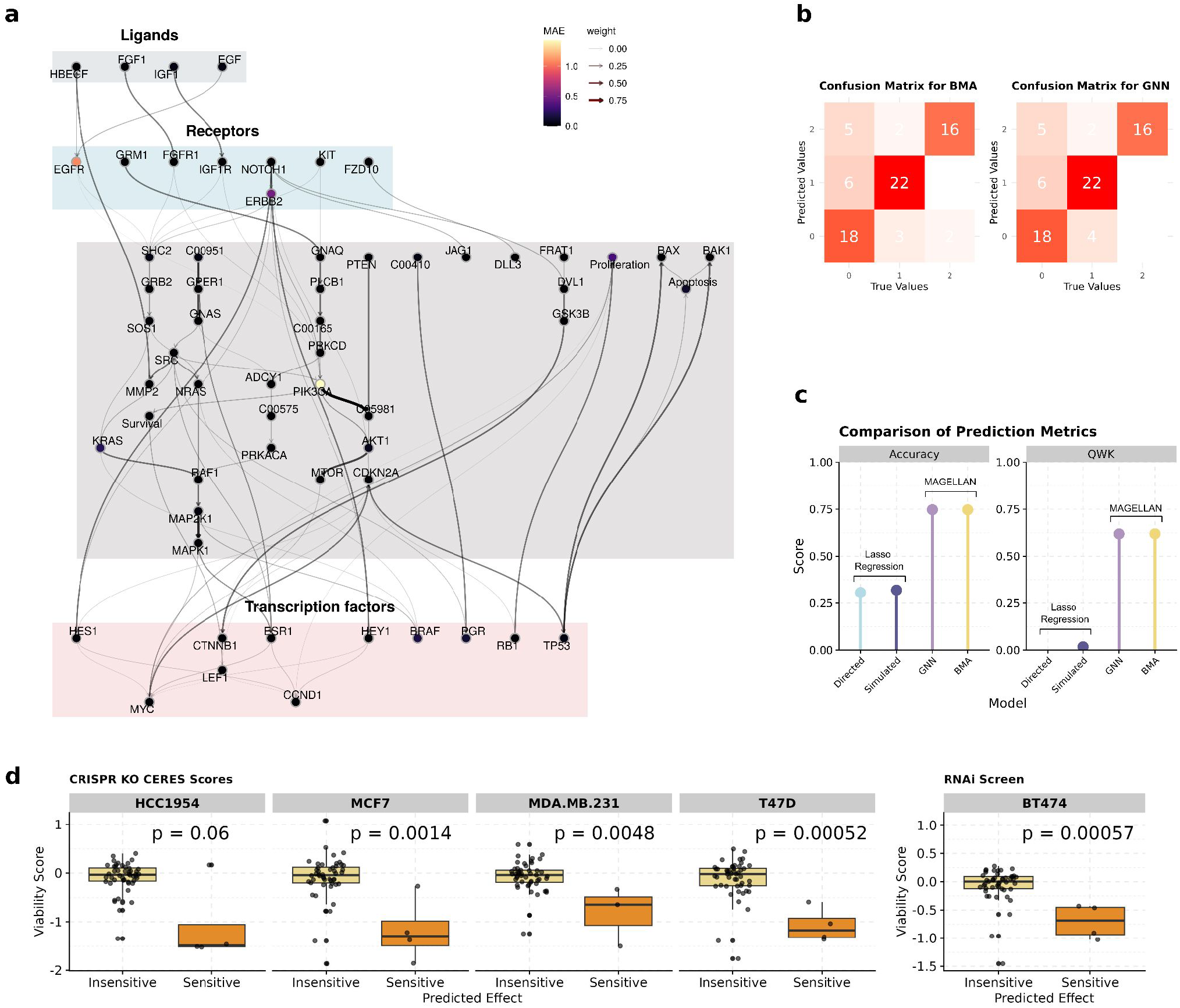
Building a computational model of breast cancer signalling using MAGELLAN. **A)** Network visualization depicting model fitting and error distribution. Nodes are coloured according to their Mean Absolute Error (MAE). Edge colour and intensity represent the fitted edge weights. **B)** Confusion matrix comparing the performance of MAGELLAN against a GNN with identical architecture. **C)** Assessment of model fit, showing accuracy and Quadratic Weighted Kappa for different methods of fitting the specification to the network. **D)** Box plots showing the gene effect (y axis, the lower the value the more sensitive the perturbation) compared to the predicted sensitivity of the perturbation according to the model (x axis). Additive scores below 0 (representing synergistic combinations) are coloured orange and additive scores 0 and above (representing insensitive combinations) are coloured pale yellow. P values are shown as calculated by the Mann-Whitney test.

We then used genetic perturbation data from DepMap (accessed from https://depmap.org) to test the model on unseen data (see Methods). CRISPR KO essentiality data^40^ was available for all but one of the cell lines tested, so for the remaining cell line (BT474) we studied RNAi essentiality^41^ on genes within the model. No data was available for the non-transformed cell line MCF10A. Sensitivity was calculated by initialising the model with cell line mutations and then simulating the models with an added perturbation. Sensitivity is calculated by the difference in predicted proliferation values between gene knockouts and untreated controls (see Methods). From this we find that the breast cancer model correctly predicts a significant decrease in viability in 4 out of 6 breast cancer cell lines (Figure 4D, Mann-Whitney, p < 0.05). Of the remaining 2 cell lines, MDA-MB-468 was not predicted to grow in conditions without perturbation, so no sensitivity could be calculated. HCC1954 had a decrease in viability of knockouts predicted to be sensitive, but with a more marginal significance (p < 0.1) compared to the rest. Overall, these results demonstrate that our MAGELLAN-optimised breast cancer model captures key genetic dependencies identified CRISPR KO and RNAi data in the majority of cell lines tested.

## Discussion

Here, we introduce MAGELLAN, a novel method for creating explainable, executable models based on qualitative rules. Importantly, this method relies on a coarse-grained, rule-based model construction rather than directly using continuous data. This allows contrasting data-formats to be integrated seamlessly together by the user, to identify emergent behaviours that arise from prior-knowledge and their own hypotheses. This was designed for transparency, accessibility and interoperability of data-input that we have illustrated by using a literature curated specification from varied provenance. Furthermore, the semi-automated nature of the fitting process allows for gaps in the knowledge to be defined and refined, directing research to the most pertinent experiments to bridge these gaps. This process can be used to help inform and minimise the number of experiments required to fit a model (a process known as active learning)^42^. This represents a powerful tool for bridging fundamental biology using sophisticated computational tools with clinically accessible utilisation.

We used MAGELLAN to re-parameterise a previously published model, showing comparable quality of fitting to training data. In this manner we illustrate that MAGELLAN effectively approximates the manually curated training process, which involved domain experts meticulously refining model parameters over several months. Importantly, MAGELLAN can do this in a fraction of the time, democratising the process of constructing sophisticated biological models to scientists without extensive computational background. We also show that MAGELLAN scales to model sizes previously unfeasible to manually trained qualitative models, showing its utility in scaling-up to large-datasets. To illustrate its utilities in a real-world example, we predict growth of breast cancer cell lines in response to various perturbations and validate on unseen data taken from DepMap. These case studies show the generalisability of MAGELLAN across diverse contexts to create interpretable biological models. Previous authors have noted that future models must distinguish imperfect and uncertain aspects of training data^42^. We have illustrated that MAGELLAN can perform satisfactorily in synthetic contexts with purposely misleading experiments in the specification. This underlines MAGELLANs ability to recover meaningful dynamics, even when the training data contains contradictions or uncertainty, inherent in biology.

Other researchers have combined machine learning with mechanistic modelling to automatically generate models from data. The algorithm cSTAR was designed by Rukhlenko *et al*. (2022)^43^ for studying and controlling cellular phenotypes using mechanistic networks generated from ‘omics data. DCell incorporates Visible Neural Networks (VNN) to couple the inner workings of machine learning to real biological ontologies, creating predictions with underlying biological mechanisms^44^. Similarly, the approach introduced by Kim *et al*. (2024)^45^ introduces a ‘grey box’ framework that combines mechanistic (“white box”) and machine learning (“black box”) modelling approaches to optimize and interpret molecular regulatory networks. This is similar to our approach in that it effectively utilises the interface between mechanistic modelling and machine learning to understand signalling networks. However, the authors note the need for drug-specific parameter tuning for optimal fitting. Our approach circumvents this limitation by using a relatively simple and standardised neural network architecture that adapts to drug-specific dynamics without requiring extensive parameter adjustments. Furthermore, through our use of qualitative networks^21^ we provide a greater number of discrete values, allowing for the capture of more complex biology than a Boolean approach. With MAGELLAN we build on this previous work, making the interface between machine learning and mechanistic modelling more generalisable to diverse contexts.

Another tool, COSMOS, uses ‘omics data expressed in the form of kinase and transcription factor activities to generate mechanistic models using network-level causal reasoning^46,47^. Similarly, SignalingProfiler 2.0 represents a tool that can explore context-specific signalling networks by integrating genomic and proteomic data with the prior knowledge-causal networks^48^. Both represent invaluable tools for mechanistic understanding of genotype to phenotype that can be used by specialist biologists understanding fundamental biological processes. However, both tools are only accessible to computational biologists with understanding of programming languages R and Python. Our integration with the BMA GUI allows wet-lab biologists to easily interrogate and modify models generated by MAGELLAN. Additionally, by framing the simulation as a propagation through a neural network, our approach is scalable to large network sizes, opening up future possibilities for integration with new ‘omics modalities and emerging data formats.

Our approach incorporates neural networks to optimise the model to fit a specification. Other authors use machine-learning models as more direct predictors of drug response. For example, PERCEPTION is a model that predicts cancer drug response using single cell data using transfer learning^12^. The transformer-based model, PROPHET, predicts cellular phenotype outcomes by learning the relationships between cellular state, treatments and phenotypic readouts across extensive experimental contexts^13^. Also, machine learning-based multi-omics has been shown to predict breast cancer cells’ sensitivity to 90 drugs^8^. However, our approach is distinct from typical machine learning models, which often focus on classification or learning hidden representations of complex data using extensive ‘deep learning’ architectures. Instead, our method explicitly emulates a process simulation, with layers representing timesteps through state-space. This results in a much simpler architecture, improving interpretability and addressing common challenges in understanding model outputs. The simplicity of the design also means that our model can be fitted with far less data, as demonstrated by the fitting of our breast cancer model to simplistic cell behaviours. This contrasts our approach starkly with more data-hungry machine learning models, that often require multi-omics and single cell resolution^49–53^ for good predictive accuracy, and which do not provide a mechanistic explanation for their predictions. Complex neural network topologies have been shown to provide rich information by these studies, but at the cost of interpretability. This abstraction makes it harder to trace predictions to the data or identify artifacts of fitting or overfitting. Our architecture is grounded in established biological knowledge. By constraining the fitting architecture to biologically relevant features, we minimize the risk of overfitting to spurious patterns, ensuring predictions are both robust, generalisable and mechanistically meaningful.

Compared to previous approaches, what sets MAGELLAN apart is its design to be efficient, user-centric, and scalable, with a Graphical User Interface (GUI) that enables researchers to interact with the resultant model intuitively. The efficiency of our method accelerates model generation and simplifies integration with diverse datasets. The GUI and the transparent rules enable peer scrutiny of both the input and the output of the results, unlike ‘black box’ or even ‘grey box’ machine learning methods.

Currently, we illustrate MAGELLAN using published models and integration with KEGG pathways. However, from our real-world examples, we see lower quality of fitting than illustrated with our synthetic benchmarks, illustrating the caveats of using real world data, fraught with noise and inaccuracies. The development of comprehensive biomedical knowledge graphs like iKraph present further opportunities for development with more structured, and high-confidence inputs that could reduce ambiguity in fitting. Such resources can enable further integration of AI into both the model architecture, facilitating more scalable, automated and literature-informed mechanistic models^54^.

Our assessment of models on drug screens shows that MAGELLAN-derived models tend to be conservative in their predictions. However, our benchmarking experiments offer insight into this behaviour, showing that increasing the number of measurements per experiment enables the model to better account for noise in both the training data and the network itself. This strongly suggests that high-quality, deep ‘omics data could be highly valuable for model training, as it allows thousands of data points to be captured in a single experiment. For future studies, incorporating such ‘omics data for training will significantly enhance both the depth and complexity of the resulting models.

The broader implication of this method is that it can effectively bridge the gap between data-driven methods and mechanistic understanding. This can open avenues for more transparent and peer-scrutinized computational modelling in clinical and research contexts. Only by democratising the advances in computational modelling and artificial intelligence, making them accessible to scientists with limited computational expertise, can we streamline the interface between experimentalists, computational biologists and clinicians. Tools, such as MAGELLAN, that facilitate this feedback between inter-disciplinary fields are vital for advancing our mechanistic understanding of intra- and inter-cellular signalling, ultimately facilitating future therapeutic breakthroughs.

## Methods

### Optimising qualitative networks using message passing graph-neural networks

MAGELLAN constructs and optimises executable, mechanistically interpretable biological network models. It begins with a user-defined signed, directed network, where nodes represent biological entities (e.g., genes, proteins, metabolites) and edges indicate their regulatory relationships (e.g., activation, inhibition).

MAGELLAN builds the model by fitting the weights of these edges to match a user-defined specification. This specification consists of a set of rules, often expressed as ‘if x, expect y’ statements, that describe the desired behaviours of the model. These rules are derived from quantitative biological observations, such as experimental data showing how the system responds to specific perturbations. The initial, ‘if x’ part of the rule is used to initialize the model, and the ‘expect y’ part is compared to the model’s simulated output. Deviations between these expectations and the simulated outcomes are used to train the edge weights, iteratively refining the model until it accurately reproduces the specified behaviours. This results in an executable model that is mechanistically interpretable, providing insights into the underlying biological processes.

To optimize the weights of the network, the model’s architecture is transformed into a message passing Graph Neural Network (GNN), allowing for efficient weight adjustment through gradient-based methods.

### Translating a Biological Prior-Knowledge Network to a Graph Neural Network (GNN) representation

Nodes in the biological network are directly related to nodes in the neural network for optimisation. These layers correspond to timesteps with each layer having as many nodes as the original biological network. The edges between layers reflect the edges within the original biological network, such that the architecture of the neural network is determined by prior knowledge.

We represent a prior-knowledge biological signalling network as a neural network by structuring a multi-layer graph that maintains the directed, causal relationships between nodes across layers. Each layer of this network captures a timestep of the simulated biological system with each node within a layer corresponding to a biological species in the original network. Edges between layers allow information to propagate in a temporally sequential manner according to the relationships in the prior-knowledge signalling network.

#### Biological network representation

The inputted network is a directed biological prior-knowledge graph, *G* = (*V,E*), where:

- *V* ={*v*_1_, *v*_2_, *v*_3_, …*v*_*n*_} represents a set of nodes (e.g., proteins, genes or chemicals),
- *E* ={*v*_*i*_, *v*_*j*_| *v*_*i*_ → *v*_*j*_} is the set of directed edges indicating activation or inhibition, according to prior knowledge.

#### Layered neural network representation

Let *L* denote the total number of layers in the network. For each node *v*_*i*_ ∈ *V* in the original network *G*, we created *L* layers with identical connections to represent L time-steps. We denote *v*_*i,l*_, *l =*1, …, *L* as the i-th node on the l-th layer. This results in an expanded set of vertices:

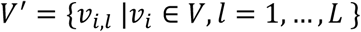

Where *v*_*i,l*_ represents the node *v*_*i*_ in layer *l*.

#### Inter-layer edges

For each prior-knowledge edge (*v*_*i*_, *v*_*j*_) ∈ *E* in the original model network *G*, we create directed and acyclic inter-layer edges between the corresponding nodes across layers in the new layered graph. Specifically, for each layer *l =* 1, …, *L* − 1, create a directed edge from *v*_*i,l*_ to *v*_*i,l*+1_:

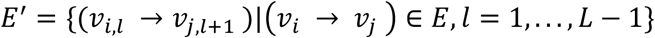

This results in the expanded, multi-layered graph *G*′.

### Propagation of regulatory signalling through layered network

The construction of *G*′ ensures that the regulatory signals propagate in a directed manner through successive layers, like time-steps. This simulates the temporal sequence of interactions as signals from one time step (layer *l*) to the next (layer *l* + 1):

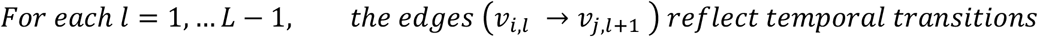

### Using MAGELLAN to fit the network to the specification

By structuring the network in this manner, *G*′ is also analogous to a neural network, where each layer represents a distinct stage in the forward propagation of information. Nodes in each layer receive “input” from their predecessors in the previous layer, while “output” edges convey the effects to nodes in the subsequent layer.

This layered approach allows us to simulate signal progression and attenuation, akin to neural network forward propagation, where edges represent synaptic connections carrying the influence from one “neuron” to another across layers (time-steps). The final layer *L* should then capture the downstream effects of the initial signalling inputs through all intermediate layers, providing insight into the cascade-like behaviour of biological signalling networks in a format that can integrate with neural network frameworks for optimisation to a qualitative specification.

For optimisation, we utilise the HardTanh activation function between layers (time steps)^55^. This enforces value bounds and constrains the node values at each time step to a fixed range (a user specified hyper parameter, e.g., 0 to 2), ensuring the updated values at each time step do not exceed the predetermined limits.

Perturbations are expressed as their own nodes (termed ‘dummy nodes’), with edges to the perturbed nodes representing the sign of the perturbations. Let *d* be a dummy node corresponding to node *D*. The relationship between the dummy node *d* and the original node *D* is defined as:

1. Add a self-loop to the dummy node (to ensure its value is maintained through all layers):

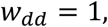
2. Connect the dummy node *d* to the original node *D* with the following logic:

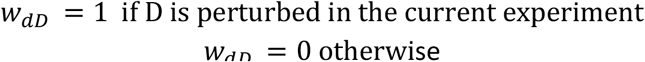
3. Assign the dummy node d fixed value *h*_*d*_, representing the perturbation value.

This ensures the incorporation of distinct perturbations across experiments, such that the topology of the neural network does not have to be modified through training rounds.

Furthermore, we introduce various edge masks to expunge unwanted perturbed edges temporarily. These masks include:

1. Edge mask *M*. ∈*R*^|*E*|^: its element equals to 0 if the corresponding edge needs to be “turned off” under the current specification experiment, e.g., self-loops for nodes that are perturbed in other specifications, as well as edges between perturbed nodes in the current specification and their parent nodes. Turning off these edges will ensure only the perturbed nodes in the current specification remain constant.
2. Perturbation mask *M*_*p*_ ∈*R*^|*E*|^: its element equals to 1 if the corresponding edges are self-loops of perturbed nodes across all specifications as well as the edges between nodes with only inhibitor parents and their dummy node parents. This ensures the weights of these edges is always 1. Note that unlike *M*_*E*_ which is different under different experimental settings, *M*_*p*_ is the same across all specifications, as *M*_*E*_ and edge index alone can turn off edges when needed.

In addition, we use an edge normalisation vector *s* ∈*R*^|*E*|^. For its *i*-th edge that connects a parent node *u* and a child node *v*, the normalisation factor is calculated by the total number of parents, *Pa*_(*v*)_, of node *v* to mimic a weighted average:

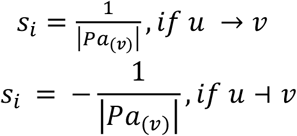

We also allow the use of a simple sum in which case:

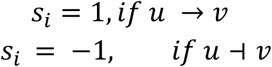

We implemented this neural network architecture to model and simulate biological signalling networks based on the specification defined previously. For classification tasks, the final node outputs are normalized using the Softmax activation function. Edge weights were initialized uniformly and trained as model parameters during execution. We used AdamW for parameter optimisation.

### Benchmarking MAGELLAN using synthetic networks

In order to assess how robust our method is to different conditions, we performed benchmarks for the effect of network size, specification size, the fraction of the specification which is incorrect (noise) and the effect of the detail per experiment on the ability to remove spurious edges while preserving correct edges (Supplementary Figure 1).

To perform these benchmarks, we use the breast cancer network derived with our method, to preserve the structure inherent in biological networks and which results from the application of the shortest path methods described in the supplementary methods section. In all cases below the metrics are calculated on a 10% holdback of the specification. Further, in all cases we use a synthetic specification, generated by choosing a random number of perturbations from within a range, and random number of measurements, and then measuring the ground truth by simulation from an initial state of all nodes at 0. To ensure that the specification has similar properties to those we have previously developed from real-world data, we bias the measured nodes toward terminally downstream nodes (e.g. phenotypes), and we bias perturbations toward either parent-less nodes (analogous to external factors such as EGF) or high-degree nodes, as we observe that perturbations in real-world experiments are often made with drugs that target such proteins. In all cases we used 30 experiment specifications unless otherwise stated.

### Benchmarking different network sizes

To generate larger networks with a similar structure to the breast cancer network, we expand the breast cancer network by iteratively duplicating randomly selected nodes, preserving most of their incoming and outgoing edges but randomly modifying a subset. In this way we can increase the size of the network while maintaining similar *e*.*g*. degree distribution and proportions of activating and inhibiting edges. To test the effect of network size on model fit, we compare the fit for different sizes with the number of experiments in the specification fixed at 25% of the nodes in the network.

### Benchmarking different specification sizes

We increase the size of the specification relative to the network, while keeping the number of perturbations and measurements per experiment within the same range.

### Benchmarking noisy and error-prone specifications

To assess the impact of noise on the method, we change the measured value of a fraction of the measured nodes per experiment.

### Benchmarking removal of spurious edges

We add *n* spurious edges to the network after the generation of a ground-truth specification. We then assess the removal of these edges while preserving true edges. We find that the method is extremely conservative, given that this requires sufficient information not only to predict the correct weights to produce the correct outputs, but to be able to distinguish between redundant pathways. However, with a large specification (100 experiments), we find that this does scale as one increases the number of measured nodes per experiment. We further find that the number of true edges removed remains very low, even for a small number of measured nodes.

### Benchmarking MAGELLAN against Lasso Regression

To benchmark the performance of the GNN approach, we implemented an alternative method based on an iterative Lasso regression model for learning edge weights. This approach treats the learning of edge weights as an optimization task with Lasso. The optimization was solved using CVXPY, a high-level modelling framework designed for convex optimization.

By enforcing sparsity through Lasso, the iterative Lasso approach identifies the most relevant connections between nodes while eliminating weaker or redundant edges. Once the edge weights were determined, they were used to model and predict experimental responses using the BMA tool.

### Re-parameterising a previously published NSCLC computational model to its specification

To model cell lines, we utilized mutational data from Van der Meer *et al*.^26^ as of 04/01/2023, as specified in the methods section of Clarke *et. al*. (2025)^24^. To train the network, we used the same specification as in Clarke *et. al*. (2025)^24^ with no hold-back set. To identify effective treatments, we modulated druggable nodes in both manually trained and MAGELLAN networks by either inactivating them (setting the target function to a minimum) or activating them (setting the target function to a maximum), both individually and in pairwise combinations. To generate a predictive measure for cell growth in both models, we calculated the difference between the values of the nodes representing proliferation and death.

For combination therapy, drug interactions were simulated and analysed using a metric called Additivity (as in Clarke *et al*. (2025)^24^), which parallels the experimentally determined Higher than Single Agent (HSA) effect from Nair *et al*. (2022)^27^. The additive score quantifies the overall survival of treated cells by measuring the difference between proliferation and death. To statistically compare the distributions of HSA values across combinations predicted to be sensitive or insensitive, we performed an independent t-test. This test evaluates whether there is a significant difference in mean HSA values between the sensitivity groups.

### Breast cancer network generation based on KEGG

We created Breast cancer KEGG pathway networks using the R package BMAlayout, with the “breast cancer” (HSA05224) and “estrogen signalling” pathways (HSA04915) serving as core inputs. We merged these KEGG pathway networks to form the basis of the model. To account for curated redundancy within KEGG, several edges were added manually, derived from KEGG or primary literature.

### Phenotype Association Layer Integration

We included associations with key phenotypes, including apoptosis, proliferation, and survival by adding relevant edges derived from the database SIGNOR^42^. We assigned interaction signs that correspond to their known regulatory roles (e.g., “activation,” “inhibition”) so that their predicted levels could be simulated in the model.

### Creating a literature-derived qualitative specification for the breast cancer computational model

We manually curated a specification defining the desired behaviour of the breast cancer computational model from a set of 10 publications describing the response of various breast cancer cell lines to different perturbations and conditions. We also integrated this with experiments from the Genomics of Drug Sensitivity in Cancer (GDSC) database.

For the GDSC-based specifications, we compiled drug viability data and Z-scores for each drug were used to rank cell lines based on their sensitivity to various compounds. The sensitivity data was filtered to retain only drugs with a Z-score greater than 1, indicating insensitivity of a cell line to the drug or below −1, indicating sensitivity of a cell line to a drug. Only drugs with consistent effects across each drug-target pair within the cell lines were kept. The complete specification for the breast cancer model is outlined in Supplementary Table S4.

### Layout Generation and Node Classification

Each pruned network was prepared for visual modelling by generating spatial layouts and classifying nodes. We used the pathwayLayout function from the R library [biotubemapR](https://github.com/CharlieBarker/biotubemapR) to define a spatial arrangement of nodes that has a biological basis. This applies protein family-based clustering separately using the Sugiyama layout algorithm for visually coherent grouping of ligands, receptors, and phenotypes.

### Breast cancer computational model assessment using integrated DepMap data

We assessed the ability of models derived from MAGELLAN with our KEGG-based BRCA network after training on the entire literature-derived specification.

For each model, we predicted simulated gene knockouts as either ‘Killing’ or ‘No effect’ for each network specification. This was done by evaluating perturbations by identifying the stable state of the network for proliferation as outlined in the ‘Simulation of the model with BioModelAnalyser’ section below. For each cell line, the model was initialised with the annotated genetics of the cell line according to CellModelPassports^24^.

To have a single value to predict cell growth the value of the ‘Proliferation’ node at stable state. If the perturbation renders the value of the ‘Proliferation’ node below that of the value of ‘Proliferation’ at the stable state of model initialised with the genetics of the cell line without a perturbation, we consider that perturbation to be killing the cell line in question, reflecting a cytostatic effect of the perturbation.

We compared these simulated results to experimental values from DepMap (https://depmap.org/portal) for CRISPR (DepMap Public 24Q2+Score, Chronos) and RNAi (Achilles+DRIVE+Marcotte, DEMETER2). ROC curves were computed to evaluate the ability of z-score ranked genes to classify samples as ‘Killing’ or ‘No effect’ as per the simulations.

### Simulation of the computational model with BioModelAnalyzer

The derived automated network is represented mathematically as a discrete qualitative network. This abstracts each gene, protein or process as a node with multiple finite values of the range (0-2). This was built, tested using the open source webtool, BioModelAnalyzer (BMA, https://biomodelanalyzer.org) and associated command line tools (https://github.com/hallba/BioModelAnalyzer).

Nodes in the network represent biological factors such as proteins or subunits of proteins. Input nodes represent growth factors. Output nodes are representing phenotypes such as apoptosis and proliferation. Regulatory interactions (activating (→) and inhibitory (⊣)) are abstracted as the edges between nodes. The activity levels of a node respond to the upstream levels of the regulators of that node, determined by a mathematical function (termed target function) for each node integrating the activities of the upstream regulators. This determines the level of the target node at the next time-step, with its activity changing by a maximum of 1 unit per time step. The target function is the weighted average of the activating upstream regulators, negated by the average of the inhibiting regulators, (*avg(pos)-avg(neg)*) or simply the sum of *(pos)-(neg)* according to user specification.

These target function weights are trained by MAGELLAN (see above) and are available for the optimised models in Supplementary Table S1 and S4.

## Code Availability

Code is available at https://github.com/jfisher-lab/MAGELLAN

## Author Contributions

Matthew A. Clarke: Conceptualisation, Methodology, Software, Validation, Writing - Original Draft; Charlie G. Barker: Conceptualisation, Methodology, Software, Validation, Formal analysis, Data Curation, Writing - Original Draft; Yuxin Sun: Conceptualisation, Methodology, Software, Data Curation, Writing - Original Draft; Theodoros I. Roumeliotis: Investigation, Resources, Data Curation, Writing - Review and Editing; Jyoti S. Choudhary: Writing - Review and Editing, Supervision, Project Administration, Funding Acquisition; Jasmin Fisher: Conceptualisation, Writing - Original Draft, Supervision, Project Administration, Funding Acquisition.

## Acknowledgements

The authors wish to thank Nir Piterman and Helena Coggan for useful discussions. J.F acknowledges funding from Cancer Research UK (C17918/A28870) and National Institute for Health Research University College London Hospitals Biomedical Research Centre. J.S.C. and T.I.R acknowledge funding from the Wellcome Trust (223745/Z/21/Z), BBSRC (BB/Y004477/1) and from the ICR.

## Supplemental Information Index

**Supplementary Table S1** – **Excel file containing network, optimised edges and derived target functions for the NSCLC computational model**. Original target functions, and their citations and rationale can be found in Clarke *et al*. (2025)^24^. Notes on the syntax of the target functions are available from https://biomodelanalyzer.org/. A number in the Target Function column represents nodes that are set to a constant value.

**Supplementary Table S2** – **Excel file containing specification and fitting for the NSCLC computational model**. For each experiment, the table shows the cell line used, the constraints applied to the model and the expected outcome, based on the experimental result. A value of 0 indicates a loss-of-function mutation conferring no activity and for 1 basal activity, 2 elevated activity and 3 strong activation.

**Supplementary Table S3** – **Excel file containing network, optimised edges and derived target functions for the breast cancer computational model**. Each row specifies a regulatory relationship between a source node (*from*) and a target node (*to*). The *Type* column indicates whether the interaction is activating or inhibiting, while the *weight* column shows the weighting of the node optimised by MAGELLAN.

**Supplementary Table S4** – **Excel file containing specification and fitting for breast cancer computational model**. For each experiment, the table shows the cell line used, the constraints applied to the model and the expected outcome, based on the experimental result. A value of 0 indicates a loss-of-function mutation conferring no activity, for 1 basal activity and 2 elevated activity in the specified measured proteins.

**Supplementary Table S5** – **List of mutations used to model each breast cancer cell line**. For every cell line, the constrained nodes and their assigned activity values are provided, representing known oncogenic mutations. A value of 0 indicates a loss-of-function mutation or deletion resulting in no activity, 1 indicates basal (normal) activity, and 2 indicates a gain-of-function mutation or amplification associated with elevated activity. The specific mutations and their functional annotations were obtained from Cell Model Passports (Van der Meer, 2019^26^).

## References

1. Sever, R. & Brugge, J. S. Signal transduction in cancer. Cold Spring Harb. Perspect. Med. 5, a006098 (2015).

2. Buchler, N. E., Gerland, U. & Hwa, T. On schemes of combinatorial transcription logic. Proc. Natl. Acad. Sci. U. S. A. 100, 5136–5141 (2003).

3. Andersen, M. E., Yang, R. S. H., French, C. T., Chubb, L. S. & Dennison, J. E. Molecular circuits, biological switches, and nonlinear dose-response relationships. Environ. Health Perspect. 110, 971–978 (2002).

4. Edwards, A. M. et al. Too many roads not taken. Nature 470, 163–165 (2011).

5. Gates, A. J., Gysi, D. M., Kellis, M. & Barabási, A.-L. A wealth of discovery built on the Human Genome Project — by the numbers. Nature 590, 212–215 (2021).

6. Garrido-Rodriguez, M. et al. Evaluating signaling pathway inference from kinase-substrate interactions and phosphoproteomics data. 2024.10.21.619348 Preprint at 10.1101/2024.10.21.619348 (2024).

7. Dugourd, A. et al. Causal integration of multi-omics data with prior knowledge to generate mechanistic hypotheses. Mol. Syst. Biol. 17, e9730 (2021).

8. Sun, R. et al. Proteomic Dynamics of Breast Cancer Cell Lines Identifies Potential Therapeutic Protein Targets. Mol. Cell. Proteomics 22, (2023).

9. Burtscher, M. L. et al. Network integration of thermal proteome profiling with multi-omics data decodes PARP inhibition. Mol. Syst. Biol. 20, 458–474 (2024).

10. Jones, I. et al. Characterization of proteome-size scaling by integrative omics reveals mechanisms of proliferation control in cancer. Sci. Adv. 9, eadd0636 (2023).

11. Barker, C. G. et al. Identification of phenotype-specific networks from paired gene expression–cell shape imaging data. Genome Res. 32, 750–765 (2022).

12. Sinha, S. et al. PERCEPTION predicts patient response and resistance to treatment using single-cell transcriptomics of their tumors. Nat. Cancer 1–15 (2024) doi:10.1038/s43018-024-00756-7.

13. Ji, Y. et al. Scalable and universal prediction of cellular phenotypes. 2024.08.12.607533 Preprint at 10.1101/2024.08.12.607533 (2024).

14. Clarke, M. A., Woodhouse, S., Piterman, N., Hall, B. A. & Fisher, J. Using State Space Exploration to Determine How Gene Regulatory Networks Constrain Mutation Order in Cancer Evolution. in Automated Reasoning for Systems Biology and Medicine (eds. Liò, P. & Zuliani, P.) 133–153 (Springer International Publishing, Cham, 2019). doi:10.1007/978-3-030-17297-8_5.

15. Howell, R. et al. Localized immune surveillance of primary melanoma in the skin deciphered through executable modeling. Sci. Adv. 9, eadd1992 (2023).

16. Howell, R. et al. Executable network of SARS-CoV-2-host interaction predicts drug combination treatments. Npj Digit. Med. 5, 1–13 (2022).

17. Kreuzaler, P. et al. Heterogeneity of Myc expression in breast cancer exposes pharmacological vulnerabilities revealed through executable mechanistic modeling. Proc. Natl. Acad. Sci. U. S. A. 116, 22399–22408 (2019).

18. Talarmain, L. et al. HOXA9 has the hallmarks of a biological switch with implications in blood cancers. Nat. Commun. 13, 5829 (2022).

19. Moignard, V. et al. Decoding the regulatory network of early blood development from single-cell gene expression measurements. Nat. Biotechnol. 33, 269–276 (2015).

20. Shorthouse, D. et al. Exploring the role of stromal osmoregulation in cancer and disease using executable modelling. Nat. Commun. 9, 3011 (2018).

21. Schaub, M. A., Henzinger, T. A. & Fisher, J. Qualitative networks: a symbolic approach to analyze biological signaling networks. BMC Syst. Biol. 1, 4 (2007).

22. Kauffman, S. Homeostasis and Differentiation in Random Genetic Control Networks. Nature 224, 177–178 (1969).

23. Clarke, M. A. & Fisher, J. Executable cancer models: successes and challenges. Nat. Rev. Cancer 20, 343–354 (2020).

24. Clarke, M. A. et al. Predicting Personalised Therapeutic Combinations in Non-Small Cell Lung Cancer Using In Silico Modelling. 2025.01.07.631497 Preprint at 10.1101/2025.01.07.631497 (2025).

25. Fisher, J. & Henzinger, T. A. Executable cell biology. Nat. Biotechnol. 25, 1239–1249 (2007).

26. van der Meer, D. et al. Cell Model Passports-a hub for clinical, genetic and functional datasets of preclinical cancer models. Nucleic Acids Res. 47, D923–D929 (2019).

27. Nair, N. U. et al. A landscape of response to drug combinations in non-small cell lung cancer. Nat. Commun. 14, 3830 (2023).

28. Kanehisa, M. & Goto, S. KEGG: kyoto encyclopedia of genes and genomes. Nucleic Acids Res. 28, 27–30 (2000).

29. Tian, J.-M., Ran, B., Zhang, C.-L., Yan, D.-M. & Li, X.-H. Estrogen and progesterone promote breast cancer cell proliferation by inducing cyclin G1 expression. Braz. J. Med. Biol. Res. Rev. Bras. Pesqui. Medicas E Biol. 51, 1–7 (2018).

30. Ahmad, S., Singh, N. & Glazer, R. I. Role of AKT1 in 17beta-estradiol- and insulin-like growth factor I (IGF-I)-dependent proliferation and prevention of apoptosis in MCF-7 breast carcinoma cells. Biochem. Pharmacol. 58, 425–430 (1999).

31. Behera, M. A. et al. Progesterone stimulates mitochondrial activity with subsequent inhibition of apoptosis in MCF-10A benign breast epithelial cells. Am. J. Physiol. Endocrinol. Metab. 297, E1089–1096 (2009).

32. Alkhalaf, M., El-Mowafy, A. & Karam, S. Growth inhibition of MCF-7 human breast cancer cells by progesterone is associated with cell differentiation and phosphorylation of Akt protein. Eur. J. Cancer Prev. Off. J. Eur. Cancer Prev. Organ. ECP 11, 481–488 (2002).

33. Moerkens, M., Zhang, Y., Wester, L., van de Water, B. & Meerman, J. H. N. Epidermal growth factor receptor signalling in human breast cancer cells operates parallel to estrogen receptor α signalling and results in tamoxifen insensitive proliferation. BMC Cancer 14, 283 (2014).

34. Sipila, P. E. et al. Prolonged tamoxifen exposure selects a breast cancer cell clone that is stable in vitro and in vivo. Eur. J. Cancer Oxf. Engl. 1990 29A, 2138–2144 (1993).

35. Liu, C.-Y. et al. Tamoxifen induces apoptosis through cancerous inhibitor of protein phosphatase 2A-dependent phospho-Akt inactivation in estrogen receptor-negative human breast cancer cells. Breast Cancer Res. BCR 16, 431 (2014).

36. Li, W. et al. Tamoxifen promotes apoptosis and inhibits invasion in estrogen-positive breast cancer MCF-7 cells. Mol. Med. Rep. 16, 478–484 (2017).

37. Hayat, A. et al. Low HER2 expression in normal breast epithelium enables dedifferentiation and malignant transformation via chromatin opening. Dis. Model. Mech. 16, dmm049894 (2023).

38. Muthuswamy, S. K., Li, D., Lelievre, S., Bissell, M. J. & Brugge, J. S. ErbB2, but not ErbB1, reinitiates proliferation and induces luminal repopulation in epithelial acini. Nat. Cell Biol. 3, 785–792 (2001).

39. Yue, W. et al. Activation of the MAPK pathway enhances sensitivity of MCF-7 breast cancer cells to the mitogenic effect of estradiol. Endocrinology 143, 3221–3229 (2002).

40. Meyers, R. M. et al. Computational correction of copy-number effect improves specificity of CRISPR-Cas9 essentiality screens in cancer cells. Nat. Genet. 49, 1779–1784 (2017).

41. Tsherniak, A. et al. Defining a Cancer Dependency Map. Cell 170, 564-576.e16 (2017).

42. Tejada-Lapuerta, A. et al. Causal machine learning for single-cell genomics. Nat. Genet. 57, 797–808 (2025).

43. Rukhlenko, O. S. et al. Control of cell state transitions. Nature 609, 975–985 (2022).

44. Ma, J. et al. Using deep learning to model the hierarchical structure and function of a cell. Nat. Methods 15, 290–298 (2018).

45. Kim, Y. et al. A gray box framework that optimizes a white box logical model using a black box optimizer for simulating cellular responses to perturbations. Cell Rep. Methods 4, 100773 (2024).

46. Dugourd, A. et al. Modeling causal signal propagation in multi-omic factor space with COSMOS. 2024.07.15.603538 Preprint at 10.1101/2024.07.15.603538 (2024).

47. Liu, A. et al. From expression footprints to causal pathways: contextualizing large signaling networks with CARNIVAL. Npj Syst. Biol. Appl. 5, 1–10 (2019).

48. Venafra, V., Sacco, F. & Perfetto, L. SignalingProfiler 2.0 a network-based approach to bridge multi-omics data to phenotypic hallmarks. NPJ Syst. Biol. Appl. 10, 95 (2024).

49. Su, G. et al. Inferring gene regulatory networks by hypergraph generative model. Cell Rep. Methods 101026 (2025) doi:10.1016/j.crmeth.2025.101026.

50. Li, C. et al. Benchmarking AI Models for <em>In Silico</em> Gene Perturbation of Cells. bioRxiv 2024.12.20.629581 (2025) doi:10.1101/2024.12.20.629581.

51. Shi, X., Ramathal, C. & Dezso, Z. INSIGHT: In Silico Drug Screening Platform using Interpretable Deep Learning Network. bioRxiv 2025.04.21.649855 (2025) doi:10.1101/2025.04.21.649855.

52. Theodoris, C. V. et al. Transfer learning enables predictions in network biology. Nature 618, 616–624 (2023).

53. Roohani, Y., Huang, K. & Leskovec, J. Predicting transcriptional outcomes of novel multigene perturbations with GEARS. Nat. Biotechnol. 42, 927–935 (2024).

54. Zhang, Y. et al. A comprehensive large-scale biomedical knowledge graph for AI-powered data-driven biomedical research. Nat. Mach. Intell. 7, 602–614 (2025).

55. Courbariaux, M., Hubara, I., Soudry, D., El-Yaniv, R. & Bengio, Y. Binarized Neural Networks: Training Deep Neural Networks with Weights and Activations Constrained to +1 or −1. Preprint at 10.48550/arXiv.1602.02830 (2016).

