## Supplementary Material for "MAGELLAN: Automated Generation of Interpretable Computational Models for Biological Reasoning"

### Supplementary Figures

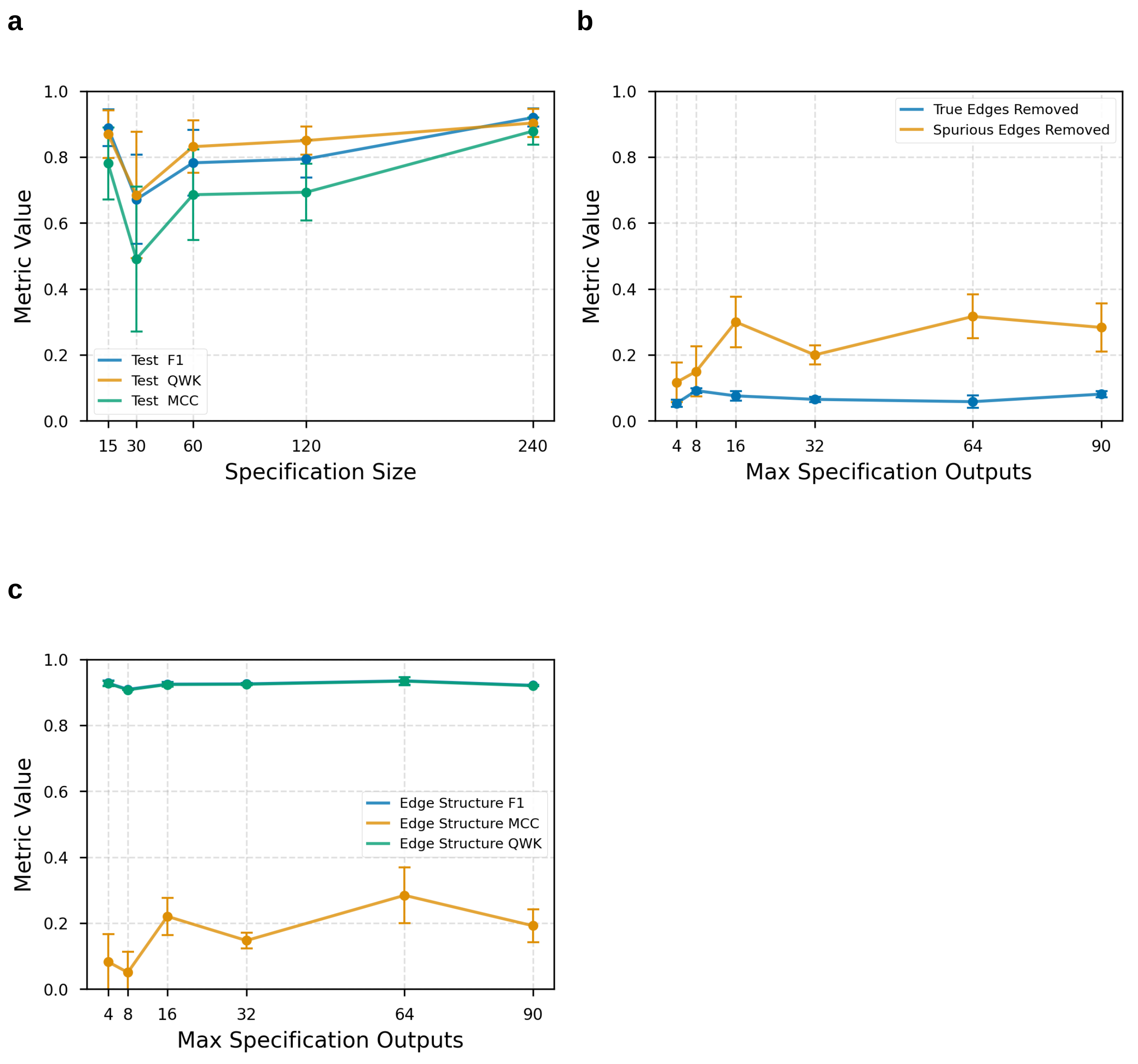


**Figure S1. Benchmarking of breast cancer computational model with synthetic specification. A)** Effect of specification size (x axis) on the quality of fitting to a test set (y axis). **B)** Proportion of edges removed (y axis) with varying number of measurements per experiment in the specification. Random edges are shown in orange and true edges are shown in blue. **C)** Impact of the number of nodes measured per experiment (x axis) on the overall quality of fitting (y axis).

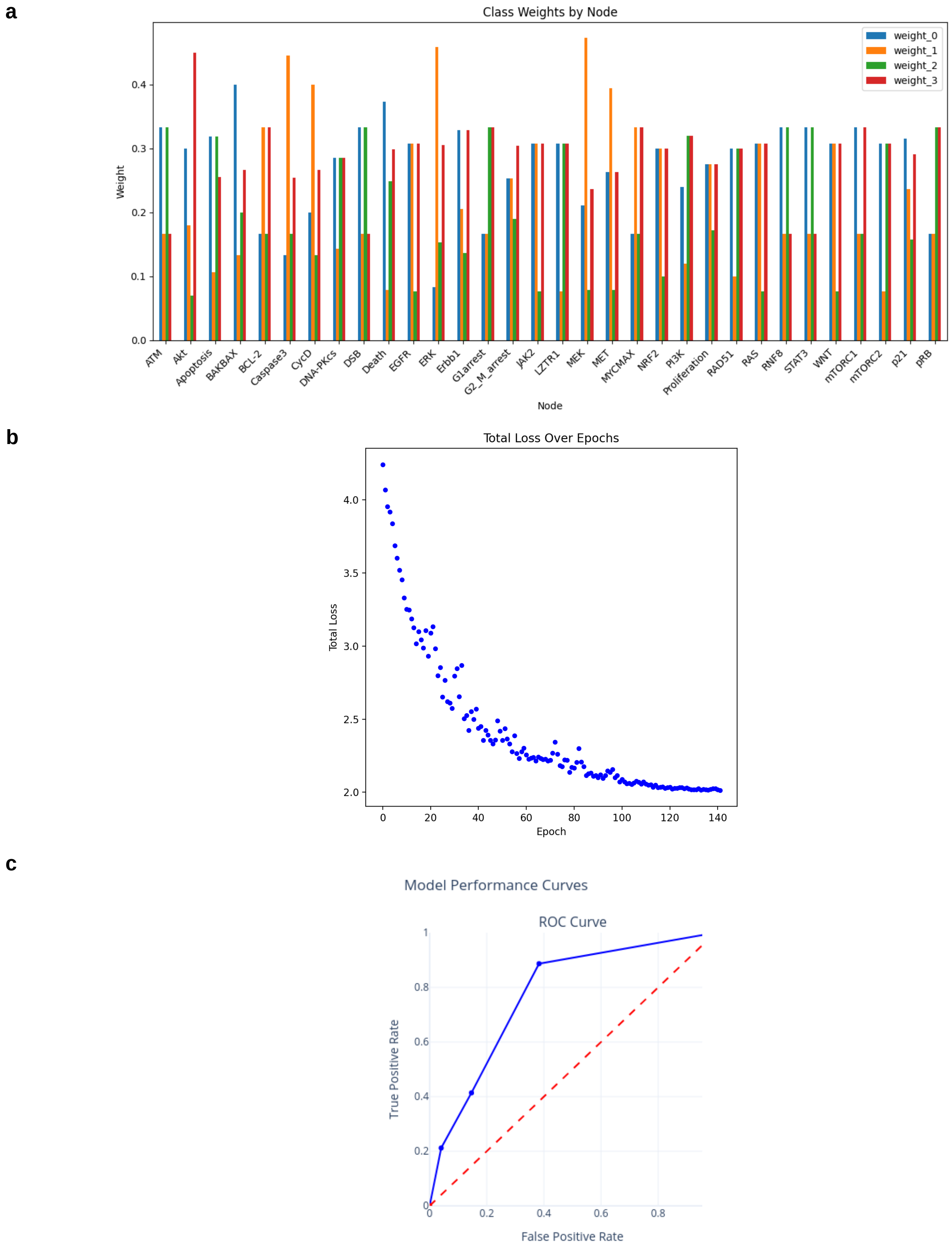


**Figure S2. Results of NSCLC computational model fitting using MAGELLAN. A)** Node-specific class weights computed according to how frequently a class appears (With rarer values given more weight). The x-axis represents individual nodes, the y-axis shows the assigned weight, and color indicates the class (value). **B)** Total training loss (y axis) across epochs (x axis) of fitting, demonstrating model convergence over time with early stopping enabled. **C)** Receiver Operating Characteristic (ROC) curve showing the trade-off between true positive (y axis) and false positive rates (x axis), evaluating the classifier's performance.).

a

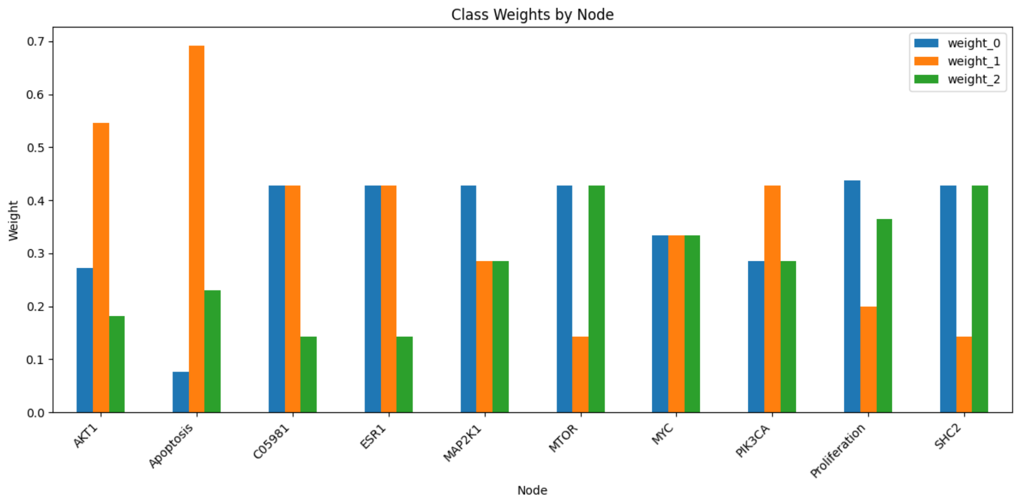


b

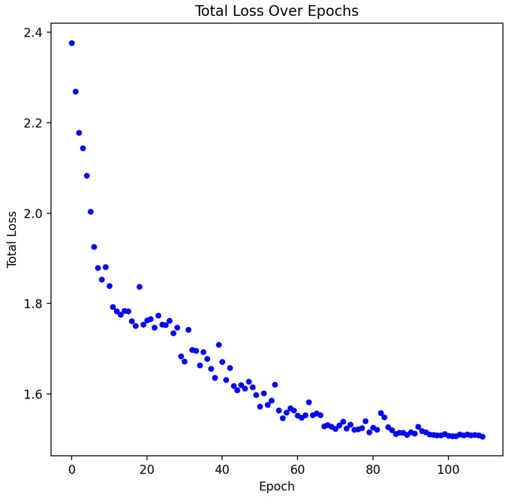


c

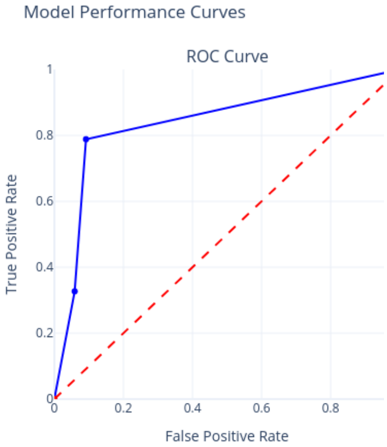


**Figure S3. Results of the breast cancer computational model fitting using MAGELLAN. A)** The x-axis represents individual nodes, the y-axis shows the assigned class weight, and color indicates the class (value). **B)** Total training loss (y axis) across epochs (x axis) of fitting, demonstrating model convergence over time with early stopping enabled. **C)** Receiver Operating Characteristic (ROC) curve showing the trade-off between true positive (y axis) and false positive rates (x axis), evaluating the classifier's performance.).

### 1 Supplementary Methods

#### 1.1 Problem definition

We search for informative interconnecting entities (genes, proteins, etc.) within a prior knowledge network $G_P$ characterised by known biological interactions. This input can be considered as an extensive directed graph, where nodes represent distinct biological entities and the edges represent the signed interactions between nodes. We aim to uncover a sparse, static graph that describes the directional interplay between the crucial biological entities specific to a cancer subtype or mechanism, and then to use known experimental data (a specification) to learn edge weights such that this graph recapitulates known results with simulated within BioModelAnalyzer (BMA) (https://biomodelanalyzer.org/) [11]. This graph, denoted as $G = (V, E)$, consists of a subset of $G_P$ in the form of nodes or vertices $V$ and the corresponding edges $E$. $V_i \in V$ represents the $i$-th node, $e_{ij} \in E$ represents edges from node $V_i$ to $V_j$. Alternatively, an edge can be represented by $V_i \rightarrow V_j$ if $V_i$ activates $V_j$, or $V_i \dashv V_j$ if $V_i$ inhibits $V_j$. Let $|S|$ denote the number of elements in a set $S$. $Pa_{(j)}$ denotes the set of parent nodes of node $V_j$, i.e., $\{i|e_{ij} \in E\}$. We further extend the definition of $Pa_{(.)}$ to $Pa^+_{(.)}$ and $Pa^-_{(.)}$ to represent the activator and inhibitor parents, respectively.

Based on their functions, the nodes $V$ are further divided into four categories:

**Genotype or mutation ($\mathcal{M}$)** Biological entities that are commonly mutated in the specific cancer type of interest, e.g. activating mutations of oncogenes such as *KRAS* [16, 25] or inactivating mutations of tumour suppressors such as *TP53* [2, 14, 19].

**Phenotype ($\mathcal{P}$)** Terminally downstream biological entities that control overall cell behaviour. For example, *BAK1* and *BAX* due to their proximity to apoptosis, and *CDK1* due to its regulation of proliferation.

**Key nodes ($\mathcal{K}$)** Measurable key nodes of interest other than those used in $\mathcal{M}$ and $\mathcal{P}$ such as *MYC* [21, 22], to be defined by the user.

**Other genes ($\mathcal{U}$)** All genes that do not fall into the above categories.

##### 1.1.1 Properties of a desired network

An optimal output network should contain the functional interactions and signalling pathways between proteins or other entities. Although the interactions are not necessarily a one-way system from genotypes to phenotypes, there should be clear paths revealing how phenotypes are determined by genotypes, possibly via key nodes. A desired network that maps genotypes to phenotypes therefore requires the following properties:

1. Should be at least a single weakly connected graph;
2. The graph should contain all the biomolecules of interest, which are defined by the user as part of the categories of nodes above;
3. Must be the minimal size to reproduce the specification so as to remain small enough to be human-readable and editable;
4. Must have a direction and a sign for all the edges;
5. Must include any direct interactions between the set of genotypes ($\mathcal{M}_i$) and phenotypes ($\mathcal{P}_j$);
6. Must include the weighted shortest paths from every genotype $\mathcal{M}_i$ to every phenotype $\mathcal{P}_j$ via key nodes $\mathcal{K}_k$, unless no paths can be discovered from existing knowledge that connects $\mathcal{M}_i$ to $\mathcal{P}_j$.

We therefore construct a problem-specific network by searching the weighted shortest paths from every genotype to every phenotype via key nodes (Figure 4). The weighted shortest paths are extracted by Dijkstra's algorithm [8].

We do not impose any constraint on path length between $\mathcal{M}_i$ and $\mathcal{P}_j$. However, within a given set of nodes of the same type (i.e. where $(i, j \in \mathcal{M}) \vee (i, j \in \mathcal{P})$), we allow at most one intermediate node. This ensures the directional flow of activity from genotype nodes toward phenotype nodes.

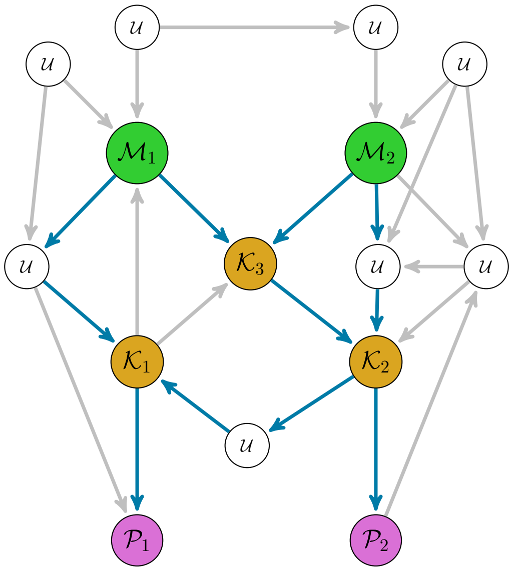


**Figure S4:** Extracting a network from directed and signed PPI interactions. Green: genotype node ($\mathcal{M}$), yellow: key node ($\mathcal{K}$, purple: phenotype node ($\mathcal{P}$). Weighted shortest paths from genotypes to phenotypes via key nodes are highlighted in blue, while uninformative interactions are coloured grey. Note that it is possible that in some cases, longer paths are preferred over shorter ones if the interactions along the longer paths have higher confidence scores such as higher number of references and thus, lower weights.

#### 1.2 Shortest paths via DEGs

##### 1.2.1 Database filtering

Our approach can use any database of intracellular interactions with a minimum set of properties. An appropriate database that suits our purpose must contain interactions that: 1) are between *human* biological entities and 2) are *directed and signed*. The sign of an interaction can be either activation or inhibition. Throughout this paper, we denote activation by $\rightarrow$ and inhibition by $\dashv$. For example, $v_1 \rightarrow v_2$ and $v_1 \dashv v_2$ mean $v_1$ activates or inhibits $v_2$, respectively.

As a proof-of-concept, we used Omnipath [4, 26, 27], a meta-database that merges protein and gene regulatory interactions from over 100 resources. Although not all interactions within Omnipath adhere to the above criteria, a notable proportion (accounting for approximately 41.1% at time of writing) constitutes valid directed and signed human PPIs. Our study focuses solely on this subset of interactions, culminating in a retained count of 21,322 directed and signed PPIs.

##### 1.2.2 Calculating edge confidence scores

Each edge $e_{ij}$ is assigned a confidence score. These scores can be evaluated from various sources including the number of references or the occurrences in various pertinent databases corresponding to the interactions. In this paper, we calculate the scores with two approaches:

1. **Reference Count**: The number of references is extracted from the “n_references” field in Omnipath.
2. **Consensus score**: While signed interactions provide valuable insights that can potentially cascade between proteins, only a limited number of databases focus on such interactions. Indeed, most PPI databases predominantly consisted of undirected or directed yet unsigned interactions. To correctly filter the interactions from the signed databases, a “consensus score” is calculated for each signed interaction. This score is based on the cumulative entries across databases, encompassing both undirected and directed variants.

The consensus score for an interaction $u \rightarrow v$ is calculated by

$$\sum_{d \in \mathcal{D}} w^{(d)} R^{(d)}_{u \rightarrow v} \quad (1)$$

where $d \in \mathcal{D}$ is a PPI database, $w^{(d)}$ is a database-specific weight, $R^{(d)}_{u\to v}$ is the number of references of interaction $u \to v$ in the database $d$. For convenience, we restrict $w^{(d)}$ to be the same for the same database types. Specifically,

$$w^{(d)} = \begin{cases} 1 & \text{if } d \text{ is undirected} \\ 2 & \text{if } d \text{ is directed} \\ 3 & \text{if } d \text{ is signed} \end{cases} \tag{2}$$

**Filtering two-edge cycles** As an initial preprocessing step, edges with scores lower than a predetermined threshold are removed to retain only high-confidence interactions within the network. These edge weights are subsequently used to filter cycles composed of two proteins $v_1 \iff v_2$. Here, the symbol $\iff$ signifies a bi-directional signal, irrespective of its sign. Within our context, in instances of opposing directions associated with varying scores, we discard the direction possessing the lower score.

##### 1.2.3 Weighted shortest paths

Once the network has been built with edges of high-confidence, we apply Dijkstra's algorithm [8] to find the weighted shortest path from each genotype node and extending to every key node, as well as from each key node to every phenotype node. This ensures the genotype-phenotype associations traverse crucial key nodes along critical signalling pathways. Intra-connections between genotype or phenotype nodes are permissible, given at most one gene to be interspersed between two nodes. The restriction of this intermittent node helps preserve a clear and unobstructed flow of activity from genotype-to-phenotype nodes within the network.

#### 1.3 Refinement and verification using BioModelAnalyzer

##### 1.3.1 Qualitative Network

One of the simplest models of discrete logic network modelling is the Boolean Network (BN) [1]. Nodes are binary, with values of 0 for "off" or 1 for "on". Nodes are connected by edges which are directed, and have signs, activation and inhibition. BNs offers a simplified yet powerful framework for depicting biological interactions. However, the binary scope confines the modelling capabilities of BNs, particularly in addressing intricate biological scenarios where entities exhibit varying levels of activity beyond binary states, such as in the case of the Myc transcription factor, which has distinct characteristics at different levels of activation [20].

In contrast, a Qualitative Network (QN) [24] capitalises on the streamlined topological structure inherent to BNs, while further extending the range of node values to discrete natural numbers. The range of nodes can be considered as granularity, indicating the number of possible states that a node can assume. In BNs, granularity is binary (0 and 1), whereas QNs have higher granularity levels. For example, the node states may be $\{0, 1, 2\}$, corresponding to no activity, low activity, and high activity. As a result, QNs are capable of modelling more intricate interactions and complex behaviours exhibited by biological systems.

A QN is a directed graph that is characterised by its nodes and edges, denoted as $V$ and $E$, respectively. Each node $V_j$ is associated with a non-negative integer state $\mathbb{Z}_*$ and a target function $f_j : \mathbb{Z}^l_* \to \mathbb{Z}_*$, mapping the states of its predecessors $\{V_i | i \in Pa_{(j)}\}$ to a scalar outcome. The default target functions are defined as

$$f_j = \begin{cases} q_j - \dfrac{\sum_{i\in Pa^+_{(j)}} x_i}{|Pa^+_{(j)}|} & \text{if } |Pa^+_{(j)}| = |Pa_{(j)}| \\ \dfrac{\sum_{i\in Pa^+_{(j)}} x_i}{|Pa^+_{(j)}|} - \dfrac{\sum_{i\in Pa^-_{(j)}} x_i}{|Pa^-_{(j)}|} & \text{otherwise} \end{cases} \tag{3}$$

where $x_i$ is the state of node $V_i$, and $q_j$ is the upper bound of node states at node $V_j$. This means that the default behaviours are characterised by the arithmetic mean of the state values exhibited by the activating parent nodes, with the corresponding value from the inhibiting parent nodes subtracted. In cases where a node solely possesses inhibiting parent nodes, the calculated average state of the activating parent nodes is substituted with the predefined upper bound of the node's state range.

However, we also allow a simple summation based target function, which we find is more robust in networks with high-degree nodes:

$$f_j = \begin{cases} q_j - \sum_{i\in Pa^+_{(j)}} x_i & \text{if } |Pa^+_{(j)}| = |Pa_{(j)}| \\ \sum_{i\in Pa^+_{(j)}} x_i - \sum_{i\in Pa^-_{(j)}} x_i & \text{otherwise} \end{cases} \tag{4}$$

To refine and verify the resulting network, we model the QN using the open-source executable biology tool BioModelAnalyzer (BMA) (https://biomodelanalyzer.org/) [11]. This tool enables us to build and test function-defined signalling networks against known cellular behaviours. By using the BMA tool, we can refine the network to ensure that it accurately reflects the known cellular behaviours. The process of refinement and verification enables us to develop a more robust and accurate network to interpret specific cancer subtypes or mechanisms based on the current literature.

##### 1.3.2 Network refinement

Pathway databases such as OmniPath [4, 26, 27] usually have an extensive collection of interactions between proteins and other biological entities. This compilation encompasses interactions that may vary in their reliability, or that could be less relevant to the problem of interest.

Hence, an additional step aimed at refining the network becomes necessary to reduce the noise of the network. One possible approach is to train the network in conjunction with BMA, in a way such that the resulting network $G$ is refined to reproduce known cellular behaviours, abstracted by a set of experiments, known as a “specification”.

The perturbation and expectation nodes across $m$ experiments are defined as $V_P = \{V_P^{(1)}, V_P^{(2)}, \cdots, V_P^{(m)}\}$ and $V_E = \{V_E^{(1)}, V_E^{(2)}, \cdots, V_E^{(m)}\}$, respectively. We represent the initial states and the pre-defined specification as $n \times m$ matrices $X$ and $Y$, respectively. Each edge $e_{ij}$ is assigned a weight $W_{ji}$, where $W$ is an $n \times n$ weight matrix. $e_{ij}$ is removed if $W_{ji} = 0$. $A$ denotes a weighted adjacency matrix, such that $A_{ji} = \pm\frac{1}{|Pa_{(j)}|}$ if $e_{ij} \in E$, and 0 otherwise (or $A_{ji} = \pm 1$ if $e_{ij} \in E$ in the case of the target function being a sum not an average). Its sign aligns with the sign of the corresponding interactions:

$$\text{sign}(A_{ji}) = \begin{cases} -1 & \text{if } V_i \dashv V_j \\ 1 & \text{if } V_i \to V_j \\ 0 & \text{otherwise} \end{cases} \tag{5}$$

To refine the network in order to reproduce known cellular behaviours in BMA, we incorporate the concept of message-passing networks, which naturally mimic BMA multi-time step behaviour with weighted default TFs by aggregating messages from their parent nodes.

In the $i$-th experiment, all nodes except the set of perturbation nodes $V_P^{(i)}$ are initialised to $0$, while $V_P^{(i)}$ is initialised to their perturbed values in this experiment. At the $t$-th iteration, every node is updated by aggregating its parents' messages from the $(t-1)$-th iteration.

$$X_j^{(t)} = \text{update}(\text{aggregate}(\{X_i^{(t-1)}, \forall i \in Pa_{(j)}\}) \tag{6}$$

$$= \sigma(\sum_{i \in Pa_{(j)}} A_{ji} W_{ji} X_i^{(t-1)}) \tag{7}$$

The message passing finishes when every node reaches a stable state and remains the same. We can then write down the expressions at the expectation nodes $V_E$, set them to the corresponding expected values in the pre-defined specification, and optimise weights $W$ over these equations. Alternatively, this is equivalent to calculating the partial derivative of $V_E$ w.r.t $V_P$. In this way, we are able to mimic multiple time steps of updates in BMA via iterations of message passing, and omit the undefined specification experiments for non-perturbation/expectation nodes.

We optimise message-passing networks with Decoupled Weight Decay Regularization (AdamW [18]) using graph neural network package PyTorch Geometric [10].

**Graph neural networks** To address the second constraint we formulate the problem in a graph neural network context, by leveraging a HardTanh [6] update function:

$$\text{HardTanh}(x, \text{min\_val}, \text{max\_val}) = \begin{cases} \text{min\_val} & \text{if } x < \text{min\_val} \\ \text{max\_val} & \text{if } x > \text{max\_val} \\ x & \text{otherwise} \end{cases} \tag{8}$$

Therefore, a single time step is executed through a dual process involving message aggregation and a HardTanh [6] activation. It is imperative to ensure that across different experiments the perturbed nodes adhere to two conditions. First, they must be correctly assigned their respective perturbations. Second, they must remain constant throughout successive time steps.

While addressing both prerequisites is readily achievable within a single experiment; by removing parent nodes and introducing self-loops for the perturbed nodes, the challenge emerges when striving to satisfy these demands across all experiments. This complexity arises due to the diverse topology required for each experiment to maintain consistent sets of perturbed nodes. In the context of a standard graph neural network, this undertaking is infeasible.

To allow us to model all experiments in a single topology, despite perturbations in effect changing the network structure, we therefore add self-loops for perturbed nodes across all experiments and expunge unwanted perturbed edges temporarily by introducing multiple masks including:

1. Edge scale: a length $|E|$ tensor vector, $s$, generated from the adjacency matrix $A_i$ with $0$ elements removed, such that for the $i$-th edge that links nodes $u$ to $v$, $s_i = \frac{1}{|Pa^+_{(v)}|}$ if $u \to v$ and $-\frac{1}{|Pa^+_{(v)}|}$ if $u \dashv v$ in the case of the default target function (Equation 3), and $s_i = 1$ if $u \to v$ and $-1$ if $u \dashv v$ in the case of the summation target function (Equation 4).

2. Edge mask $M_e$: a length $|E|$ binary tensor. Its element equals to 0 if the corresponding edge needs to be "turned off" under the current experiment, e.g. self loops for nodes that are perturbed in *other* experiments, and edges between perturbed nodes in the *current* experiment and their parent nodes. Turning off these edges will ensure only the perturbed nodes in the current experiment remain constant.

3. Perturbation mask $M_p$: a length $|E|$ binary tensor. Its element equals to 1 if the corresponding edges are self-loops of perturbed nodes across *all* experiments as well as the edges between nodes with only inhibitor parents and their "dummy" node parents. This ensures the weights of these edges is always 1. Note that unlike $M_e$ which is different under different experiment settings, $M_p$ is the same across all experiments, as $M_e$ and edge index alone can "turn off" necessary edges.

In addition, we need to add a dummy node that is indexed by 0, as adding masks to edge index is equivalent to "turning off" edges by changing the corresponding index to 0. This inevitably introduce many self loops to the node indexed by 0.

Let $x$ denote a length $|N| + 1$ tensor, where $x_i$ is the initial state of the $i - 1$-th node. Note that $x_1 = 0$, representing the dummy node indexed by 0. Let $I$ denote a $2 \times |E|$ tensor such that $I_{.,j}$ represents the node indexes for the $j$-th edge. Assume $\odot$ represents an element-wise multiplication, or, in the case where a length $m$ vector $x \odot$ a matrix $X$ with $m$ entries along one of its dimensions (e.g. the 2-nd dimension), the $\odot$ operator is equivalent to perform $x_i * X_{.,i}$ for $i = 1, \cdots, m$. The feed forward step is shown in Algorithm 1.

**Algorithm 1** Feed forward for a single experiment

```
Input: initial node states x, edge index I, edge mask M_e, perturbation mask M_p, edge scale s, maximum
node value n̂, stacked SimpleConv class with sum aggregation C
Mask edges I ← I ⊙ M_e
W ← W ⊙ (1 − M_p) + M_p
W ← W ⊙ s
for conv ∈ C do
    x' ← conv(x, I, W)
    x' ← x + HardTanh(x' − x, 0, 1)
    x ← HardTanh(x', 0, n̂)
end for
Return x
```

After a fixed set of feed-forward iterations, the predictions of the network for the nodes measured in the current experiment within the specification is compared to the ground-truth in order to calculate the loss. As these predictions are $\mathbb{R}$ but our true values are $\mathbb{Z}_*$, this is a problem of multi-label classification of ordinals. Accordingly, we choose to use Earth-Mover Distance as our loss function [23], which has been suggested as an effective metric in this class of problems [3, 7, 9]. Finally, due to the large class imbalances between nodes (e.g. proliferation is above 0 for all experiments except those in which a cytostatic effect is achieved) we can weigh the loss per node inversely with how often a result of that class is observed for that node across the entire specification (Supplementary Figure 3).

#### 1.4 Benchmarking

Benchmarks were performed on an Apple M3 Pro with 36GB of unified memory and 4.05GHz CPU.

#### 1.5 Network Generation

We generate a larger network from a base model (in this case our breast cancer model) while retaining the overall structure using a duplication-divergence approach [15]. To do this we:

1. **Select a node**
2. **Duplicate the node and associated edges**.
3. **Divergence** For each edge for this node either:
   - **Duplicate with the same properties** Duplicate the edge with the same weight and sign.
   - **Create a diverged edge.** Duplicate the edge changing the weight and/or sign.
   - **Delete edge** Remove the edge from the network.
4. **Repeat** Continue the process till a desired network size is reached.

#### 1.6 Specification Generation

In order to generate the specification for a synthetic network, for each experiment we:

1. **Choose a set of perturbed nodes.** For nodes with at least one child, we choose $n$ nodes within a user defined range, biased towards those that are either parent-less or have a high node-degree, and biased towards nodes not already used as perturbations.
2. **Choose a set of measured nodes.** For nodes with at least one parent, we choose $n$ nodes within a user defined range, biased toward nodes that are childless i.e. representing terminally downstream genes and proteins, and biased towards nodes not already used as perturbations.
3. **Generate the ground-truth values.** For each set of perturbations, we simulate the effect of perturbations in the network and fill the specification with the ground-truth values for the measured nodes.

We bias the selection of nodes to mimic those that are seen in real experimental data, as detailed in our prior work [5, 12, 13, 17].

### References


[1] Stefan Bornholdt. Boolean network models of cellular regulation: prospects and limitations. *Journal of The Royal Society Interface*, 5:S85–S94, 2008. doi: 10.1098/rsif.2008.0132.focus. URL https://royalsocietypublishing.org/doi/abs/10.1098/rsif.2008.0132.focus.

[2] Jean-Christophe Bourdon, Kenneth Fernandes, Fiona Murray-Zmijewski, Geng Liu, Alexandra Diot, Dimitris P. Xirodimas, Mark K. Saville, and David P. Lane. p53 isoforms can regulate p53 transcriptional activity. *Genes & Development*, 19(18):2122–2137, 2005. doi: 10.1101/gad.1339905. URL http://genesdev.cshlp.org/content/19/18/2122.abstract.

[3] Alberto Castaño, Pablo González, Jaime Alonso González, and Juan Jose Del Coz. Matching distributions algorithms based on the earth mover's distance for ordinal quantification. *IEEE Transactions on Neural Networks and Learning Systems*, 35(1):1050–1061, 2022.

[4] Francesco Ceccarelli, Denes Turei, Attila Gabor, and Julio Saez-Rodriguez. Bringing data from curated pathway resources to Cytoscape with OmniPath. *Bioinformatics*, 36(8):2632–2633, 12 2019. ISSN 1367-4803. doi: 10.1093/bioinformatics/btz968. URL https://doi.org/10.1093/bioinformatics/btz968.

[5] Matthew A Clarke, Charlie George Barker, Ashley Nicholls, Matt P Handler, Lisa Pickard, Amna Shah, David Walter, Etienne De Braekeleer, Udai Banerji, Jyoti Choudhary, et al. Predicting personalised therapeutic combinations in non-small cell lung cancer using in silico modelling. *bioRxiv*, pages 2025–01, 2025.

[6] Matthieu Courbariaux, Itay Hubara, Daniel Soudry, Ran El-Yaniv, and Yoshua Bengio. Binarized neural networks: Training deep neural networks with weights and activations constrained to+ 1 or-1. *arXiv preprint arXiv:1602.02830*, 2016.

[7] Giovanni Da San Martino, Wei Gao, and Fabrizio Sebastiani. Ordinal text quantification. In *Proceedings of the 39th International ACM SIGIR conference on Research and Development in Information Retrieval*, pages 937–940, 2016.

[8] E. W. Dijkstra. A note on two problems in connexion with graphs. *Numerische Mathematik*, 1(1):269–271, 1959. doi: 10.1007/BF01386390. URL https://doi.org/10.1007/BF01386390.

[9] Andrea Esuli, Fabrizio Sebastiani, and Ahmed Abbasi. Sentiment quantification. *IEEE Intell. Syst.*, 25(4): 72–75, 2010.

[10] Matthias Fey and Jan Eric Lenssen. Fast graph representation learning with pytorch geometric. *arXiv preprint arXiv:1903.02428*, 2019.

[11] Jasmin Fisher and Thomas A Henzinger. Executable cell biology. *Nature Biotechnology*, 25(11):1239–1249, 2007. doi: 10.1038/nbt1356. URL https://doi.org/10.1038/nbt1356.

[12] Rowan Howell, Matthew A Clarke, Ann-Kathrin Reuschl, Tianyi Chen, Sean Abbott-Imboden, Mervyn Singer, David M Lowe, Clare L Bennett, Benjamin Chain, Clare Jolly, et al. Executable network of sars-cov-2-host interaction predicts drug combination treatments. *NPJ Digital Medicine*, 5(1):18, 2022.

[13] Rowan Howell, James Davies, Matthew A Clarke, Anna Appios, Inês Mesquita, Yashoda Jayal, Ben Ringham-Terry, Isabel Boned Del Rio, Jasmin Fisher, and Clare L Bennett. Localized immune surveillance of primary melanoma in the skin deciphered through executable modeling. *Science Advances*, 9 (15):eadd1992, 2023.

[14] M. Isobe, B. S. Emanuel, D. Givol, M. Oren, and C. M. Croce. Localization of gene for human p53 tumour antigen to band 17p13. *Nature*, 320(6057):84–85, 1986. doi: 10.1038/320084a0. URL https://doi.org/10.1038/320084a0.

[15] Iaroslav Ispolatov, Pavel L Krapivsky, and Anton Yuryev. Duplication-divergence model of protein interaction network. *Physical Review E—Statistical, Nonlinear, and Soft Matter Physics*, 71(6):061911, 2005.

[16] Onno Kranenburg. The kras oncogene: Past, present, and future. *Biochimica et Biophysica Acta (BBA) - Reviews on Cancer*, 1756(2):81–82, 2005. ISSN 0304-419X. doi: https://doi.org/10.1016/j.bbcan.2005.10.001. URL https://www.sciencedirect.com/science/article/pii/S0304419X05000624. The KRAS Oncogene.

[17] Peter Kreuzaler, Matthew A Clarke, Elizabeth J Brown, Catherine H Wilson, Roderik M Kortlever, Nir Piterman, Trevor Littlewood, Gerard I Evan, and Jasmin Fisher. Heterogeneity of myc expression in breast cancer exposes pharmacological vulnerabilities revealed through executable mechanistic modeling. *Proceedings of the National Academy of Sciences*, 116(44):22399–22408, 2019.

[18] Ilya Loshchilov and Frank Hutter. Decoupled weight decay regularization. *arXiv preprint arXiv:1711.05101*, 2017.

[19] O W McBride, D Merry, and D Givol. The gene for human p53 cellular tumor antigen is located on chromosome 17 short arm (17p13). *Proceedings of the National Academy of Sciences*, 83(1):130–134, 1986. doi: 10.1073/pnas.83.1.130. URL https://www.pnas.org/doi/abs/10.1073/pnas.83.1.130.

[20] Daniel J Murphy, Melissa R Junttila, Laurent Pouyet, Anthony Karnezis, Ksenya Shchors, Duyen A Bui, Lamorna Brown-Swigart, Leisa Johnson, and Gerard I Evan. Distinct thresholds govern myc's biological output in vivo. *Cancer cell*, 14(6):447–457, 2008.

[21] Chadd E Nesbit, Jean M Tersak, and Edward V Prochownik. Myc oncogenes and human neoplastic disease. *Oncogene*, 18(19):3004–3016, 1999. doi: 10.1038/sj.onc.1202746. URL https://doi.org/10.1038/sj.onc.1202746.

[22] Stella Pelengaris, Mike Khan, and Gerard Evan. c-myc: more than just a matter of life and death. *Nature Reviews Cancer*, 2(10):764–776, 2002. doi: 10.1038/nrc904. URL https://doi.org/10.1038/nrc904.

[23] Yossi Rubner, Carlo Tomasi, and Leonidas J Guibas. A metric for distributions with applications to image databases. In *Sixth international conference on computer vision (IEEE Cat. No. 98CH36271)*, pages 59–66. IEEE, 1998.

[24] Marc A. Schaub, Thomas A. Henzinger, and Jasmin Fisher. Qualitative networks: a symbolic approach to analyze biological signaling networks. *BMC Systems Biology*, 1(1):4, 2007. doi: 10.1186/1752-0509-1-4. URL https://doi.org/10.1186/1752-0509-1-4.

[25] Nobuo Tsuchida, Tom Ryder, and Eiichi Ohtsubo. Nucleotide sequence of the oncogene encoding the p21 transforming protein of kirsten murine sarcoma virus. *Science*, 217(4563):937–939, 1982. doi: 10.1126/science.6287573. URL https://www.science.org/doi/abs/10.1126/science.6287573.

[26] Dénes Türei, Tamás Korcsmáros, and Julio Saez-Rodriguez. Omnipath: guidelines and gateway for literature-curated signaling pathway resources. *Nature Methods*, 13(12):966–967, 2016. doi: 10.1038/nmeth.4077. URL https://doi.org/10.1038/nmeth.4077.

[27] Dénes Türei, Alberto Valdeolivas, Lejla Gul, Nicolàs Palacio-Escat, Michal Klein, Olga Ivanova, Márton Ölbei, Attila Gábor, Fabian Theis, Dezső Módos, Tamás Korcsmáros, and Julio Saez-Rodriguez. Integrated intra- and intercellular signaling knowledge for multicellular omics analysis. *Molecular Systems Biology*, 17(3):e9923, 2021. doi: https://doi.org/10.15252/msb.20209923. URL https://www.embopress.org/doi/abs/10.15252/msb.20209923.
